# A broad exome study of the genetic architecture of asthma reveals novel patient subgroups

**DOI:** 10.1101/2020.12.10.419663

**Authors:** Sophia Cameron-Christie, Alex Mackay, Quanli Wang, Henric Olsson, Bastian Angermann, Glenda Lassi, Julia Lindgren, Michael Hühn, Yoichiro Ohne, Monica Gavala, Jingya Wang, Gundula Povysil, Sri V. V. Deevi, Graham Belfield, Inken Dillmann, Daniel Muthas, Suzanne Cohen, Simon Young, Adam Platt, Slavé Petrovski

## Abstract

**Introduction:** Asthma risk is a complex interplay between genetic susceptibility and environment. Despite many significantly-associated common variants, the contribution of rarer variants with potentially greater effect sizes has not been as extensively studied. We present an exome-based study adopting 24,576 cases and 120,530 controls to assess the contribution of rare protein-coding variants to the risk of early-onset or all-comer asthma.

**Methods:** We performed case-control analyses on three genetic units: variant-, gene- and pathway-level, using sequence data from the Scandinavian Asthma Genetic Study and UK Biobank participants with asthma. Cases were defined as all-comer asthma (n=24,576) and early-onset asthma (n=5,962). Controls were 120,530 UK Biobank participants without reported history of respiratory illness.

**Results:** Variant-level analyses identified statistically significant variants at moderate-to-common allele frequency, including protein-truncating variants in *FLG* and *IL33.* Asthma risk was significantly increased not only by individual, common *FLG* protein-truncating variants, but also among the collection of rare-to-private *FLG* protein-truncating variants (p=6.8×10^−7^). This signal was driven by early-onset asthma and did not correlate with circulating eosinophil levels. In contrast, a single splice variant in *IL33* was significantly protective (p=8.0×10^−10^), while the collection of remaining *IL33* protein-truncating variants showed no class effect (p=0.54). A pathway-based analysis identified that protein-truncating variants in loss-of-function intolerant genes were significantly enriched among individuals with asthma.

**Conclusions:** Access to the full allele frequency spectrum of protein-coding variants provides additional clarity about the potential mechanisms of action for *FLG* and *IL33.* Beyond these two significant drivers, we detected a significant enrichment of protein-truncating variants in loss-of-function intolerant genes.

## Introduction

Asthma is the 10^th^ highest cause of Disability Adjusted Life Years (DALYs) worldwide[1], affecting >300 million adults[2]. Though geographic prevalence varies[2,3], asthma remains the most common chronic condition in children[3]. With prevalence rising over time[3,4] the global burden of asthma provides a significant public health challenge. Despite the recent emergence of effective therapies, predominantly targeting elevated type 2 inflammation[5], only four novel therapy mechanisms have been FDA approved since 2000[6]. This leaves many individuals living with asthma that is poorly controlled and treated stochastically, on symptomatology rather than underlying pathobiology[7]. To successfully develop novel therapeutics, a ‘precision medicine’ approach is vital to target treatments to patient populations most likely to benefit[8].

Twin studies estimate asthma heritability at 35% in adults[9,10] and 80% in children[11,12], suggesting genetics plays a major part in disease development, particularly in early-onset asthma. Over the last two decades, the primary tool for identifying genetic asthma factors was microarray-based, genome wide association studies (GWAS). These identified and replicated dozens of common single nucleotide polymorphisms and multi-gene loci correlated to asthma risk, including the 6p21 HLA region, 17q12-21 (*ORMDL3, GSDML* and *IKZF3*)[13], and individual genes such as *IL33[14]*. Most observed odds ratios for asthma GWAS are <1.1, occasionally as high as 2 (EBI GWAS Catalogue[15]).

The contribution of rare variants to asthma risk has not been thoroughly studied. Despite numerous, pedigree-based studies of familial asthma[16,17], the only variants reported in ClinVar[18] or the Online Mendelian Inheritance in Man (OMIM[19]) are two relatively common protein-truncating variants in the gene *filaggrin* (*FLG*), each with a minor allele frequency (MAF) of 1-2% in West-European populations[20], associated with allergic phenotypes and childhood asthma[21]. To overcome a lack of power with individual rare variants, gene-unit methods such as collapsing analysis[22,23] have identified known and novel disease-associated genes.

We present the first comprehensive, exome-wide sequence analysis of a large asthma population. Our study focuses on the genetic architecture of asthma by investigating the contribution of protein-coding variation across three genomic test units: 1) variant-level, exome-wide association study (exWAS), 2) gene-based collapsing analysis, and 3) pathway-based burden analysis. We applied these methods to >24,000 participants with asthma from the UK Biobank (UKB) and an AstraZeneca clinical trial, compared to exomes from more than 120,000 participants of the UKB without asthma or respiratory conditions **(Figure 1)**. Additionally, given the high heritability of childhood asthma[11,12] early-onset cases may carry more high-risk variants, so we also extended our analyses to the early-onset (<18 years old) asthma sub-group.

**Figure 1.**
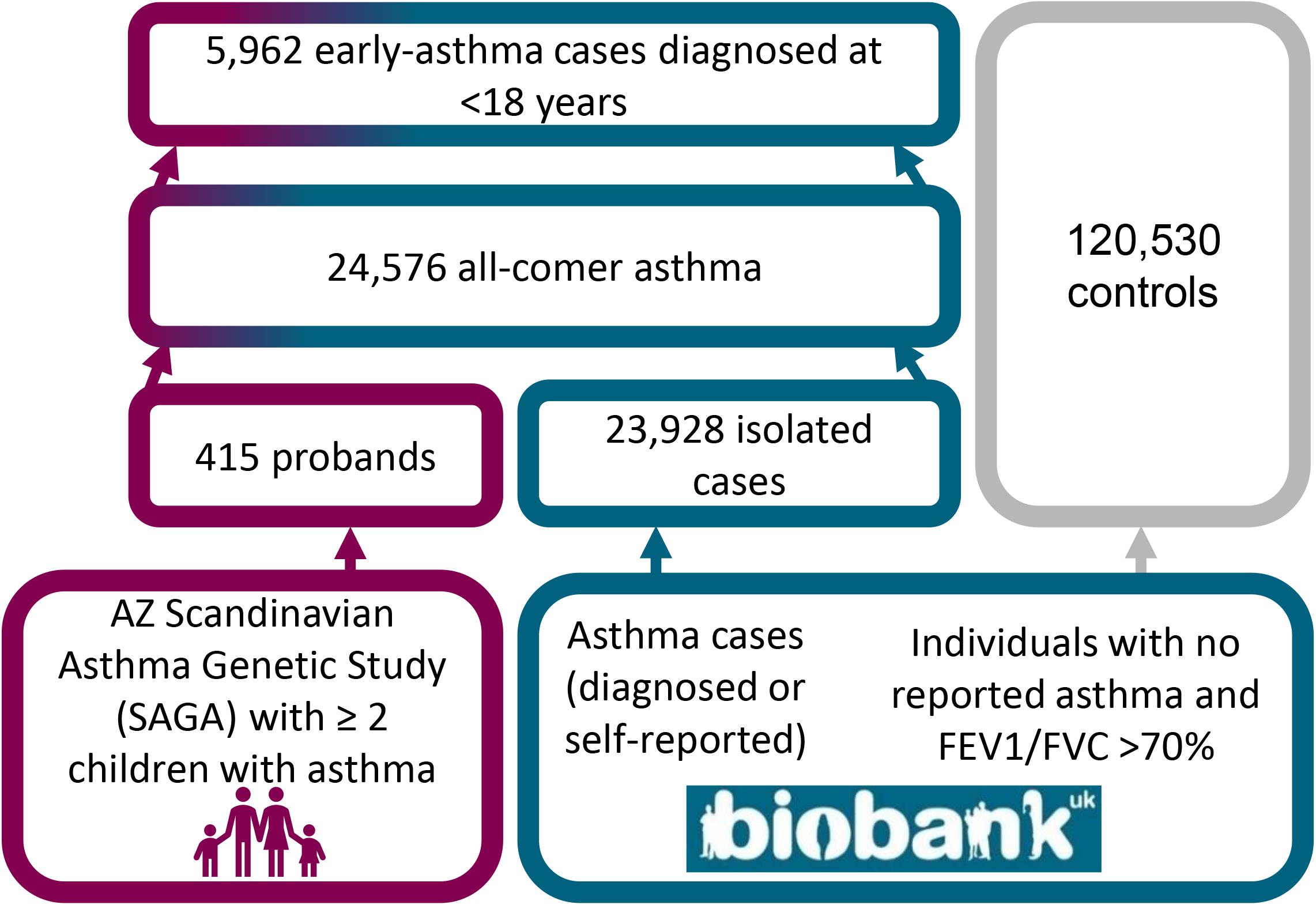
Case and control cohorts.

## Methods

### Clinical cohorts

The Scandinavian Asthma Genetic Study (SAGA) is an AstraZeneca clinical trial that ran between 1999 and 2000. Participants were recruited from 10 centres spanning Denmark, Norway and Sweden. The original study involved 532 families, of which at least two children had a clinical diagnosis of asthma (Section S1), totalling 2,392 participants. Genomes of 422 index subjects from distinct families were sequenced.

The UK Biobank is a project which recruited circa 500,000 adults aged 40-69 years in 2006-2010 from 22 centres across Britain. Participants provided DNA, biomarkers, past and ongoing medical records, and underwent further health assessments and monitoring. This resource is available to approved researchers for health-related research in the public interest (www.ukbiobank.ac.uk).

### Ethics

Ethics committees at all recruiting institutions approved the SAGA study, which was performed in accordance with the provisions of the Declaration of Helsinki as defined by the International Conference on Harmonization, Good Clinical Practice, and applicable regulatory requirements, and the policy on bioethics and human biologic samples of the trial sponsor, AstraZeneca. During the study, participants or guardians provided consent for the collection and use of DNA and clinical data for genetic research related to asthma and/or asthma-related conditions.

Ethical approval for the UK Biobank (UKB) was granted by the UK National Health Service National Research Ethics Service (Research Ethics Committee approval number 11/NW/0382). Participants provided consent for general research consistent with the UK Biobank stated purpose, to improve the prevention, diagnosis, and treatment of illness and the promotion of health throughout society.

### Case Selection

Two groups of cases were defined: ‘all asthma’ and ‘early asthma’ for cases diagnosed at <18 years of age.

SAGA families were of North-West-European ancestry. Median age at diagnosis was three years for cases. UKB cases were selected if they had an ICD diagnosis of asthma (**Supplementary Material S1**) or if they self-reported a diagnosis of asthma (Field ID 20002, code 1111). Age of UKB self-reported asthma diagnoses were 0–76 years, median 33.5 **(Figure S1)**.

After quality filtering **(Table S1)**, the all-comer asthma group comprised of 415 SAGA and 24,161 UKB participants with an asthma diagnoses. For the ‘early asthma’ group, UKB subjects were included with self-reported diagnosis at <18 years of age (UKB n=5552 and SAGA n=410).

### Control Selection

Control subjects were selected from the UKB if they lacked a self-reported or ICD10 diagnosis of asthma, or any other major respiratory illness **(Table S1)**, and their UKB records reported an FEV1/FVC>70%. From 200,603 participants with exome sequence data, this totalled 120,530 UKB controls.

### Sequencing and Bioinformatics

Genomic DNA from UKB participants underwent paired-end 75bp whole exome sequencing (WES) at Regeneron Pharmaceuticals using IDT xGen v1 capture kit and NovaSeq6000. SAGA subjects were extracted and underwent paired-end 150bp WGS at Human Longevity Inc using the NovaSeq6000 platform. For all UKB participants, >95% of CCDS had at least 10x coverage and average coverage of CCDS was 59X. For both cases and controls, alignment to GRCh38 and SNV and indel calling were performed using a custom-built Amazon Web Services (AWS) cloud compute platform running Illumina DRAGEN Bio-IT Platform Germline Pipeline v3.0.7. SNVs and indels were annotated using SnpEFF v4.3 against Ensembl Build 38.92 (Section S2).[24] SNVs and indels with MAPQ<30 were excluded from all analyses. Demographic checks of sex and predicted ancestry were performed (Section S2).

### Subject Quality Control (QC)

Exomes were excluded if they didn’t qualify a set of subject-level QC, including contamination levels, discordance between reported and genetic-predicted sex, low coverage of the target exome boundaries, and read-depth outliers **(Table S1)**. Due to the available data, this study focused on subjects of European genetic ancestry **(Table S1)**.

### exWAS (variant level)

We performed an exome-wide association study (exWAS) across all protein-coding variants seen ≥8 times in the combined case-control cohort. We studied four genetic modes of inheritance: allelic, genotypic, dominant and recessive.

### Collapsing Analysis (gene level)

In collapsing analysis, we aggregate variants fitting a given set of criteria within each gene, identified as ‘qualifying variants’ (QVs).[22] We performed 11 non-synonymous collapsing analyses, 10 dominant and one recessive model, plus an additional synonymous variant model as a negative control. In each model, for each gene, the proportion of cases is compared to the proportion of controls carrying one or more qualifying variants in that gene, except the recessive model, where a subject must have two qualifying alleles. The criteria for qualifying variants in each collapsing analysis model are in **Table S3**, all with identical QC filters **(Table S2)**.

### MegaGene (gene-set) Analysis

We applied MegaGene[25] to investigate biologically relevant gene-sets and pathways enriched for genes with qualifying variants in the case or control groups. We tested 9,341 pre-defined gene-sets for each genetic model (Supplementary material S3). Briefly, we applied a logistic regression model comparing disease status to the number of genes containing qualifying variants (qualifying genes, QGs), with three covariates: sex, number of synonymous QVs per individual per gene set (accounting for differences in neutral variation rate between gene sets), and the exome-wide tally of QVs an individual has in the tested model.

### Statistics

Calculations performed in R software (version 3.6.2).[26] For exWAS, we defined significance as the conventional p<5×10^−08^. For collapsing analysis, we used a two-sided Fisher’s exact test for P-values with a conservative, Bonferroni, study-wide significance threshold as p<2.6×10^−7^ (0.05/[(18695 genes)*(10 non-synonymous models)]). Because gene-sets have considerable overlap, we focus on gene-sets achieving Benjamini & Hochberg (FDR) adjusted p<0.01.

## Results

Gene-based collapsing analyses identified a number of highly-ranked genes of potential interest **(Table 1 and S4, Figure S3)**, including study-wide significant results for *FLG* and near-significance for *IL33*. ExWAS analyses identified 45 variants associated with asthma risk in known loci **(Table S5)**. Finally, gene-set analysis identified a previously unreported case enrichment of PTVs in loss-of-function intolerant genes **(Figure 4, Table S6)**.

**Table 1.** Top 20 gene results from non-synonymous collapsing analysis, ranked by lowest p-value

| Group | Model | n cases | Gene Name | Qual Case Freq | Qual Ctrl Freq | Enriched Direction | FET P | Odds Ratio | Odds Ratio LCI | Odds Ratio UCI |
| --- | --- | --- | --- | --- | --- | --- | --- | --- | --- | --- |
| early asthma | PTVcommon | 5962 | *FLG* | 0.1780 | 0.1168 | Case | 5.2x10<sup>-41</sup> | 1.64 | 1.53 | 1.75 |
| all asthma | PTVcommon | 24579 | *FLG* | 0.1370 | 0.1168 | Case | 2.3x10<sup>-18</sup> | 1.20 | 1.15 | 1.25 |
| all asthma | PTVcommon | 24579 | *IL33* | 0.0079 | 0.0114 | Control | 8.7x10<sup>-7</sup> | 0.69 | 0.60 | 0.81 |
| early asthma | flexnonsyn | 5962 | *CAMK1D* | 0.0050 | 0.0018 | Case | 3.6x10<sup>-6</sup> | 2.74 | 1.87 | 4.02 |
| early asthma | rec | 5962 | *CA12* | 0.0008 | 1.7E-5 | Case | 4.5x10<sup>-6</sup> | 50.58 | 9.81 | 260.78 |
| early asthma | rec | 5962 | *PALM3* | 0.0007 | 0.0000 | Case | 4.9x10<sup>-6</sup> | Inf | NA | NA |
| all asthma | raredmgmtr | 24579 | *RELB* | 0.0007 | 0.0001 | Case | 5.2x10<sup>-6</sup> | 4.91 | 2.55 | 9.43 |
| early asthma | flexnonsynmtr | 5962 | *CAMK1D* | 0.0042 | 0.0014 | Case | 6.7x10<sup>-6</sup> | 2.96 | 1.95 | 4.51 |
| early asthma | rec | 5962 | *ASCC3* | 0.0017 | 0.0002 | Case | 7.5x10<sup>-6</sup> | 7.23 | 3.51 | 14.89 |
| early asthma | raredmg | 5962 | *TGFB1* | 0.0040 | 0.0014 | Case | 1.1x10<sup>-5</sup> | 2.95 | 1.92 | 4.53 |
| early asthma | flexnonsyn | 5962 | *MRGPRX3* | 0.0099 | 0.0052 | Case | 1.1x10<sup>-5</sup> | 1.92 | 1.47 | 2.51 |
| all asthma | flexnonsyn | 24579 | *CAMK1D* | 0.0033 | 0.0018 | Case | 1.1x10<sup>-5</sup> | 1.81 | 1.41 | 2.34 |
| all asthma | raredmg | 24579 | *CNGB1* | 0.0037 | 0.0059 | Control | 1.3x10<sup>-5</sup> | 0.63 | 0.50 | 0.78 |
| all asthma | raredmg | 24579 | *TEDDM1* | 0.0000 | 0.0005 | Control | 2.1x10<sup>-5</sup> | Inf | NA | NA |
| all asthma | ptv | 24579 | *ACTRT3* | 0.0004 | 4.2E-5 | Case | 2.5x10<sup>-5</sup> | 9.81 | 3.35 | 28.71 |
| all asthma | PTVcommon | 24579 | *ACTRT3* | 0.0004 | 4.2E-5 | Case | 2.5x10<sup>-5</sup> | 9.81 | 3.35 | 28.71 |
| all asthma | flexnonsynmtr | 24579 | *CAMK1D* | 0.0027 | 0.0014 | Case | 2.7x10<sup>-5</sup> | 1.90 | 1.43 | 2.52 |
| early asthma | flexnonsyn | 5962 | *SH3BP5* | 0.001 | 0.004 | Control | 3.9x10<sup>-5</sup> | 0.25 | 0.11 | 0.56 |
| early asthma | flexdmg | 5962 | *SH3BP5* | 0 | 0.0018 | Control | 4.2x10<sup>-5</sup> | 0 | NA | NA |
| early_asthma | raredmgmtr | 5962 | *SLC35A4* | 0.002 | 0.00044 | Case | 4.6x10<sup>-5</sup> | 4.58 | 2.45 | 8.58 |

**FET** = Fisher's Exact Test, **LCI/UCI** = Lower/Upper Confidence Interval (95%). Model parameters are shown in Table S3.

### *FLG* loss-of-function variants significantly increase risk of childhood asthma

In gene-based collapsing analysis of PTVs with MAF<5% we observed a study-wide significant association between *FLG* PTVs (OMIM#135940) in early-onset (p=5.2×10^−41^, OR=1.64, 95%CI 1.53-1.75), and all asthma (p=2.3×10^−18^, OR=1.20, 95%CI 1.15-1.25), **Figure 2**; **Table 2**). Beyond the two well-known common *FLG* PTVs, we observed an additional 269 distinct *FLG* PTVs in our cohort; 115 (43%) were private, seen in only a single subject. The two most common *FLG* variants (MAF>1%)[21,27], rs61816761-G-A and rs558269137-CACTG-C, are observed among 10.6% of all cases. The remaining variants observed, each at less than 1% MAF, accounted for the remaining 3.4% of cases. In total, 3,368/24,567 cases carry at least one putative truncating variant in *FLG* (13.7%). When we restricted the PTV collapsing analysis in *FLG* to a MAF<1%, the collection of rare *FLG* PTVs remained highly significant, with a consistent odds ratio to common PTVs (p-value 6.8×10^−7^, OR=1.22, 95%CI=1.13-1.32) providing firm evidence that *FLG* PTVs as a class increase asthma risk.

**Figure 2.**
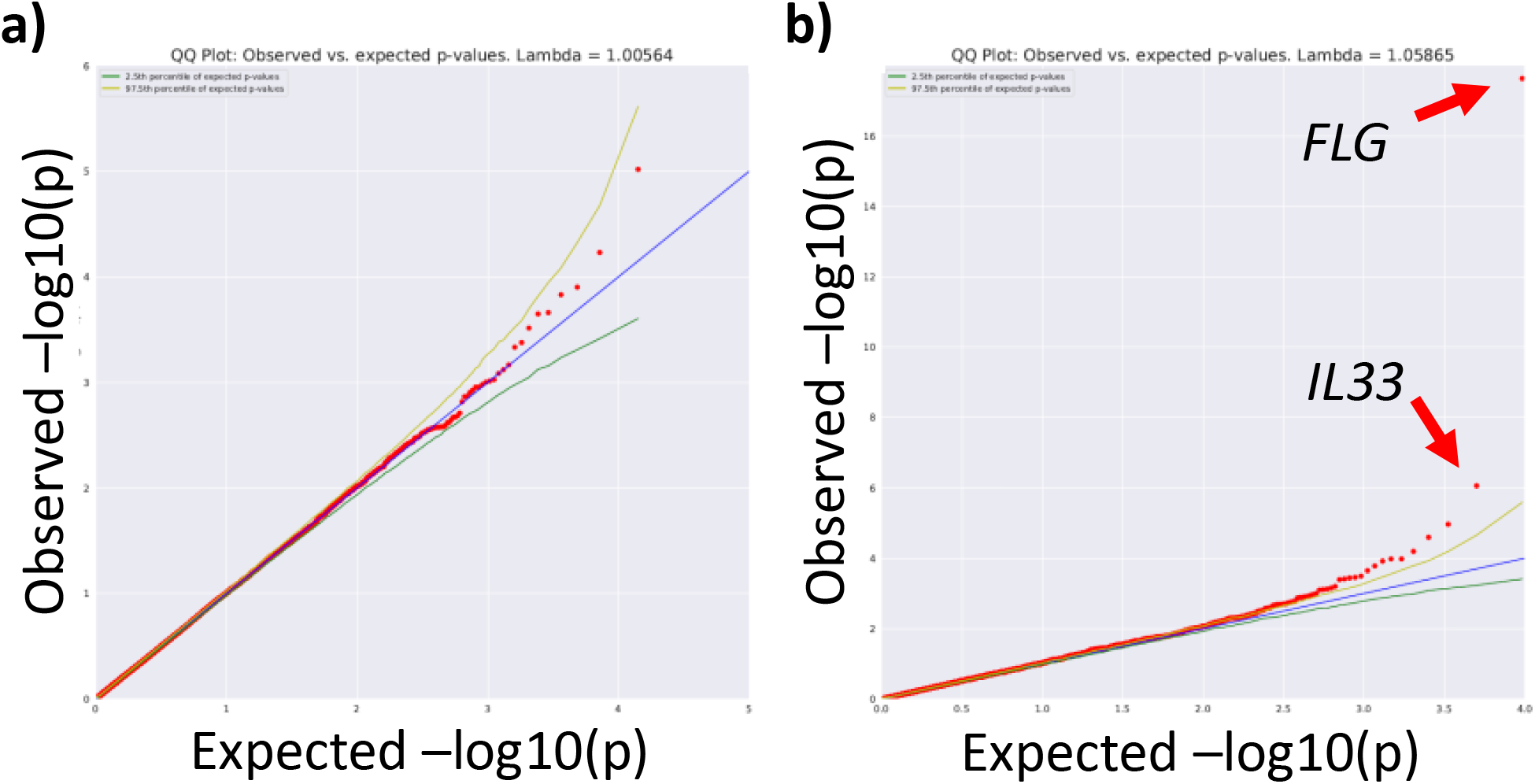
Collapsing analysis of genes in all-comer asthma shows that synonymous variants (negative control) matched the expected distribution of p-values while among the minor allele frequency (MAF) <5% protein truncating variant (PTV) collapsing model, cases are significantly enriched for PTVs in *FLG*. (A) synonymous variants, (B) PTV only, MAF<5% filter

**Table 2.**
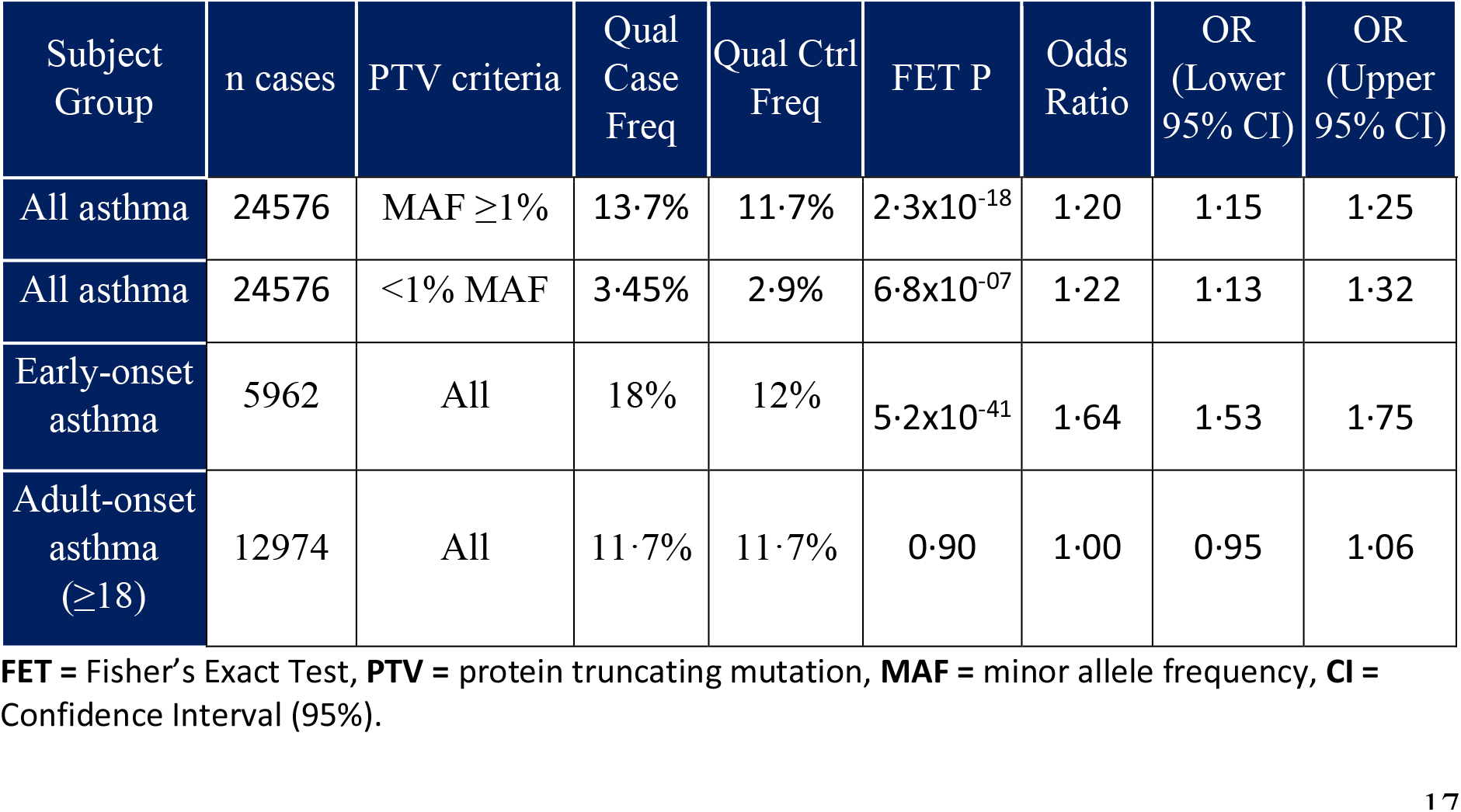
Effect of *FLG* PTV variants on asthma risk

| Subject Group | n cases | PTV criteria | Qual Case Freq | Qual Ctrl Freq | FET P | Odds Ratio | OR (Lower 95% CI) | OR (Upper 95% CI) |
| --- | --- | --- | --- | --- | --- | --- | --- | --- |
| All asthma | 24576 | MAF ≥ 1% | 13.7% | 11.7% | 2.3x10<sup>-18</sup> | 1.20 | 1.15 | 1.25 |
| All asthma | 24576 | <1% MAF | 3.45% | 2.9% | 6.8x10<sup>-07</sup> | 1.22 | 1.13 | 1.32 |
| Early-onset asthma | 5962 | All | 18% | 12% | 5.2x10<sup>-41</sup> | 1.64 | 1.53 | 1.75 |
| Adult-onset asthma (≥18) | 12974 | All | 11.7% | 11.7% | 0.90 | 1.00 | 0.95 | 1.06 |

**FET** = Fisher's Exact Test, **PTV** = protein truncating mutation, **MAF** = minor allele frequency, **CI** = Confidence Interval (95%).

To further examine the reported relationship between *FLG* PTV carriers and asthma, we stratified the UK Biobank self-declared participants into groups with asthma onset of <10 years, 10-18 years, or ≥18 years. The strongest effect of carrying a *FLG* PTV was on the youngest asthma group (p=8.5×10^−37^, OR=1.82, 95%CI 1.67-1.99), with a moderate effect at 10-18 years old (p=1.9×10^−7^, OR=1.38, 95%CI 1.22-1.55), and no measurable effect on the rate of asthma in adults (p=0.9, OR=1.00, 95%CI 0.95-1.06) **(Table 2)**.

The effect of recessive *FLG* loss, in homozygous or likely compound heterozygous PTV carriers, is even stronger (**Table 3, Figure 3)**. Biallelic carriers of *FLG* PTVs have over a 9-fold increase in asthma diagnosed before 10 years old (p=1.0×10^−53^, OR=9.2, 95%CI 7.27-11.56), a 3.5-fold increase of diagnoses between 10–18 years (p=2.3×10^−7^, OR=3.5, 95%CI 2.25-5.30), and over 1.5-fold increase of adult diagnoses (p=2.7×10^−5^, OR=1.77, 95%CI 1.36-2.27).

**Table 3.**
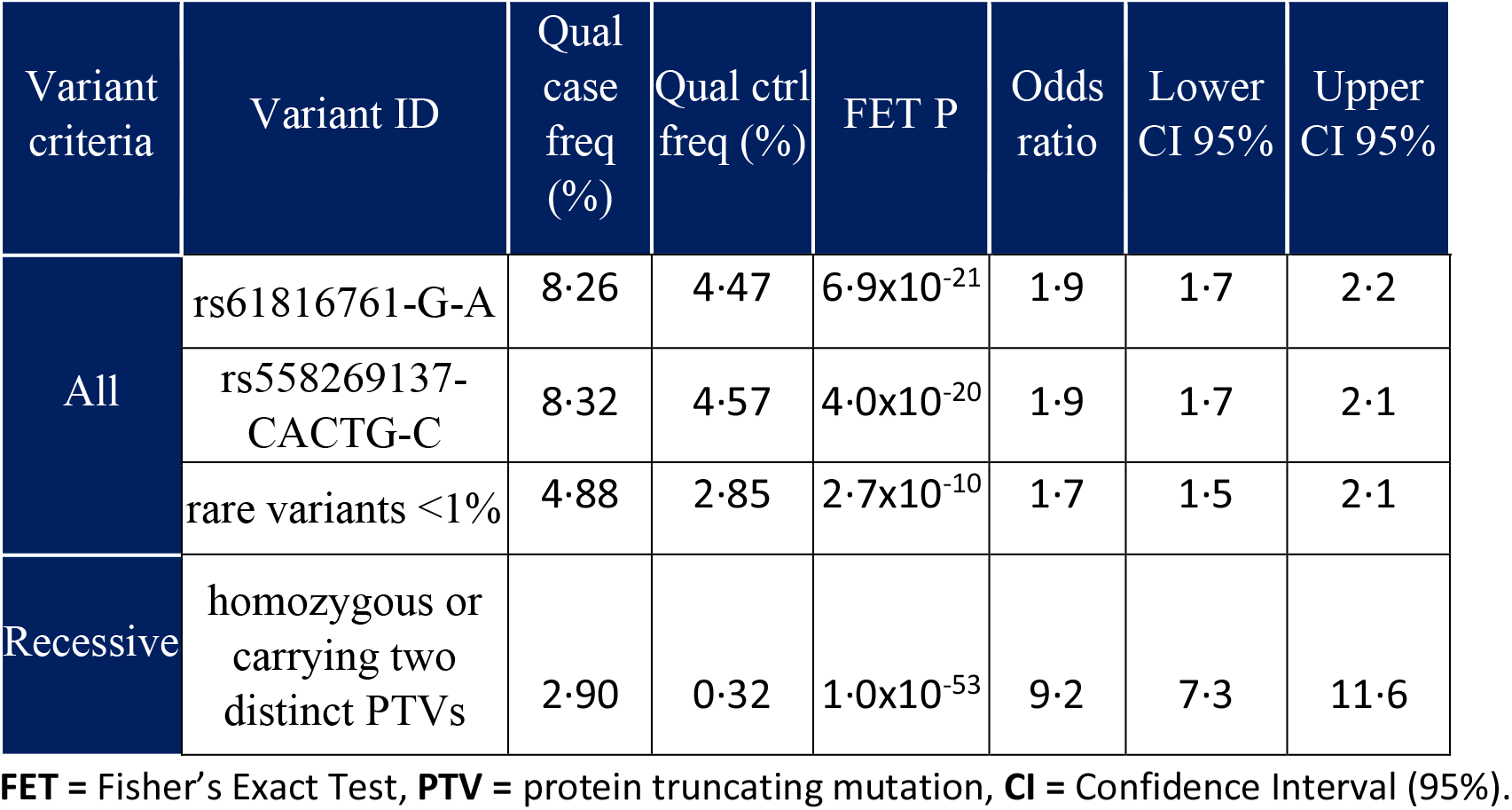
Effect of *FLG* PTV variants in <u>self-reported</u> UKB asthma cases diagnosed at <10 years old (n=3342). ‘Recessive’ includes homozygotes and putative compound heterozygotes.

| Variant criteria | Variant ID | Qual case freq (%) | Qual ctrl freq (%) | FET P | Odds ratio | Lower CI 95% | Upper CI 95% |
| --- | --- | --- | --- | --- | --- | --- | --- |
| All | rs61816761-G-A | 8.26 | 4.47 | $6.9 \times 10^{-21}$ | 1.9 | 1.7 | 2.2 |
| | rs558269137-CACTG-C | 8.32 | 4.57 | $4.0 \times 10^{-20}$ | 1.9 | 1.7 | 2.1 |
| | rare variants <1% | 4.88 | 2.85 | $2.7 \times 10^{-10}$ | 1.7 | 1.5 | 2.1 |
| Recessive | homozygous or carrying two distinct PTVs | 2.90 | 0.32 | $1.0 \times 10^{-53}$ | 9.2 | 7.3 | 11.6 |

**FET** = Fisher’s Exact Test, **PTV** = protein truncating mutation, **CI** = Confidence Interval (95%).

**Figure 3.**
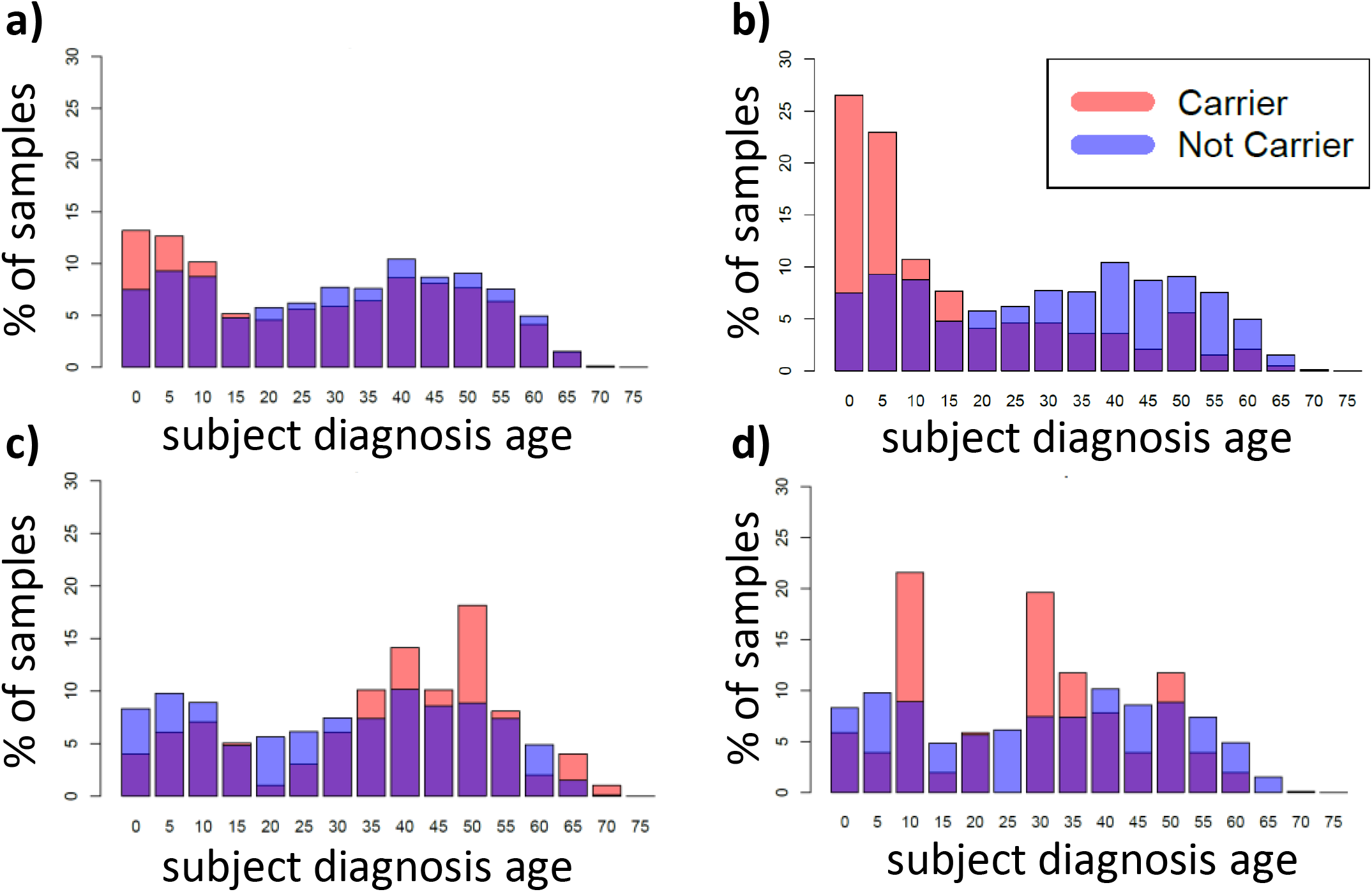
Distribution of age at asthma diagnosis in UK BioBank for **(A)** All *FLG* protein truncating variant (PTV) variant carriers (n=2511, red), versus *FLG* PTV non-carriers (n=16,015, blue), P=1.90×10^−21^. **(B)** Recessive *FLG* PTV carriers (n=196, red) versus *FLG* PTV non-carriers (n=16,015, blue), P=7.9×10^−28^. **(C)** Age at diagnosis for carriers of the *IL33* rs146597587-C protective splice-site variant (n=99, red), versus homozygous reference cases with no *IL33* PTVs (n=18,376, blue), P=0.001. **(D)** Case carriers of an alternative *IL33* PTV other than rs146597587-C (n=51, red) versus non-carriers (n=18,376, blue), P=0.46.

Because common *FLG* PTVs have been associated with increased allergy phenotypes[27], most studies investigating *FLG* and asthma posit that *FLG* perturbation and skin barrier dysfunction leads to an increase in allergy risk, resulting in allergic asthma risk. Consistent with this, asthma patients with a *FLG* PTV were more likely to self-report, or be diagnosed, with an allergic phenotype compared to *FLG* carriers with no asthma (p=1.5×10^−143^ OR=3.58, 95%CI 3.26-3.94). However, when excluding UK Biobank participants who had eczema, atopic dermatitis, psoriasis, or allergic rhinitis, among other phenotypes **(Supplementary material S1)** the relationship between *FLG* PTV carrier status and early-onset asthma remained significant (<18 y.o., p=3.4×10^−14^, OR=1.42, 95%CI 1.30-1.55).

### A protective IL-33 variant is less prevalent in early-onset asthma

A previously reported asthma-associated gene, *IL33*, (OMIM#608678) was depleted of PTVs among all asthma cases in the UK Biobank (p=8.7×10^−7^, OR=0.69, 95%CI=0.60-0.81), with greater depletion in early-onset asthma (p=4.9×10^−4^, OR=0.59, 95%CI=0.43-0.80) (**Figure 3, Table 4**). Closer review of the underlying data determined the signal was driven by a single splice variant found at a MAF of 0.36% among non-Finnish Europeans[28] (p=8.0×10^−10^, OR=0.58, 95% CI: 0.48-0.70). This replicates findings from a previous report[24] that the *IL33* rs146597587-C splice variant is depleted among individuals with asthma. That study[24] demonstrated a loss-of function phenotype when one rs146597587-C allele was present. Compared to controls, we observed that *IL33* rs146597587-C carrier rates were significantly lower in early-onset (<18 y.o.) asthmatics compared to later onset (0.39% carriers in early vs. 0.61% in late onset, Fisher’s exact p=0.001). Both groups are lower than controls (0.88%).

**Table 4.**
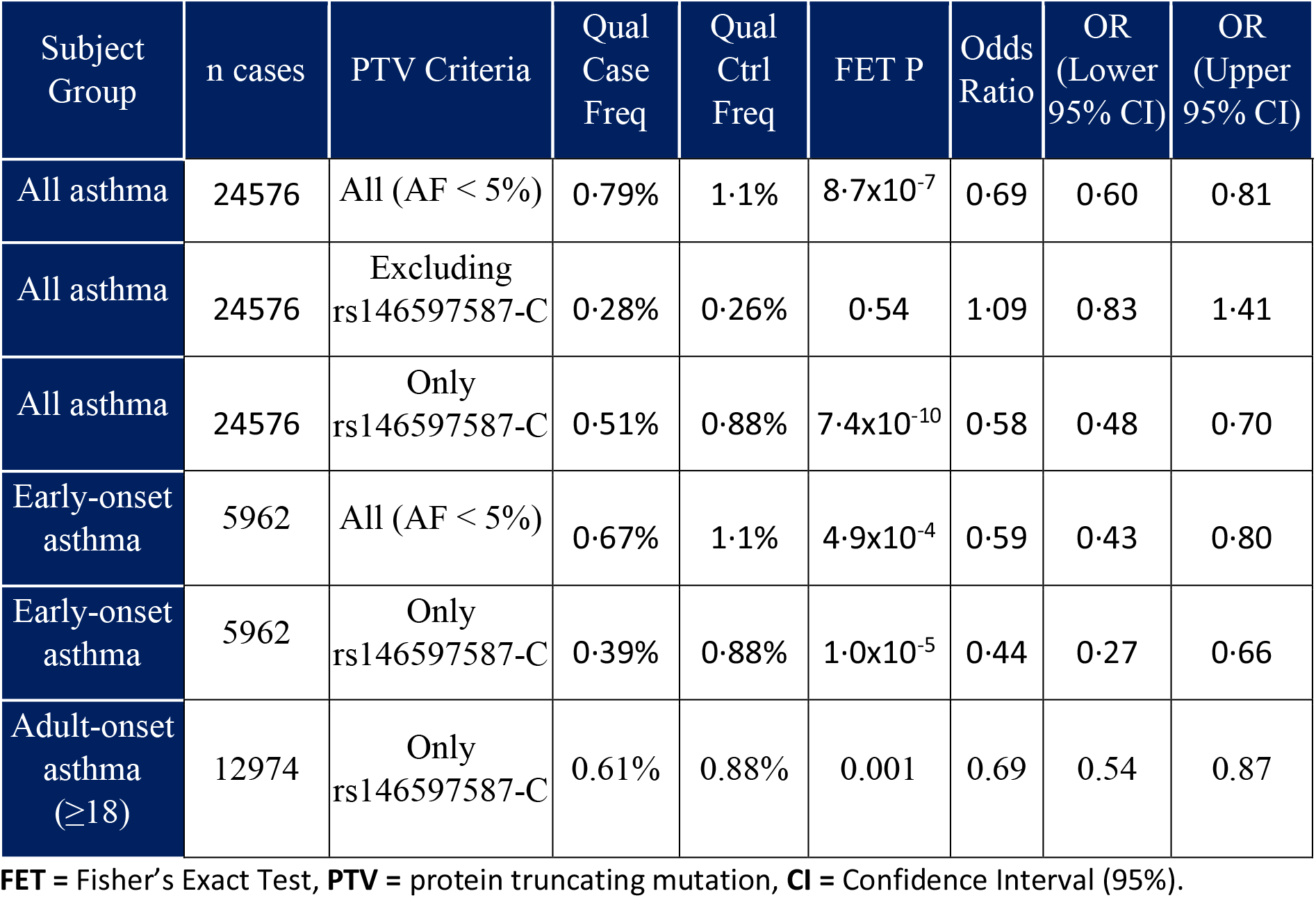
Effect of *IL33* PTV variants on asthma risk

In our cohort we identified an additional 380 individuals carrying one of 13 rare *IL33* PTVs (highest MAF=0.13%). Notably, unlike the class effect in *FLG* variants, the remaining *IL33* PTVs do not achieve a significant relationship with asthma status (p=0.54, OR=1.09, 95%CI=0.83-1.41) (**Table 4**).

### Relationship between asthma risk variants and eosinophil counts

To determine whether the carrier status of *FLG* PTVs, carriers of *IL33* splice-site rs146597587-C, or carriers of other *IL33* PTVs have a relationship with blood eosinophil counts, we examined the UKB, European participants with both exome sequence data and eosinophil levels available (n=144,570). We applied rank-based inverse normal transformation to normalise the eosinophil count distribution (UKB Field 30150, instance 0). After correcting for recruitment age and sex in linear regression, we confirmed previous reports[24] that the 1,184 carriers of the rs146597587-C *IL33* splice variant had significantly lower eosinophil counts than the 143,386 non-carriers, irrespective of disease status (p=2.3×10^−23^; β=-0.29, standard error=0.03), but no significant relationship was observed with eosinophil counts among 380 carriers of other *IL33* PTVs (p=0.10; β=-0.08, standard error=0.05), and no relationship among 17,374 carriers of *FLG* PTVs (p=0.13; β=-0.01, standard error=0.01).

### Variant Analysis

In the all-comer asthma group, 45 variants achieved p<5×10^−8^ **(Table S5)**. Thirty-two (70%) were inside the 6p21 HLA-associated locus and six (13%) were in the previously implicated 17p21 locus[13]. The remaining eight loci (19%), which include the two most common *FLG* PTVs, have all been previously reported in association with asthma, with the exception of a variant in *RPTN*, which appears to be in linkage with a *FLG* PTV. No protein-coding variants in the nearby 17p12 locus, also linked to asthma[13], achieved p<1×10^−6^. In the early asthma group, the 40 significant variants were already identified as significant in the all-asthma group, but 32 (80%) reported higher odds ratios in early asthma **(Table S6)**, consistent with previous reports[21].

### MegaGene analysis

Two gene-sets achieved FDR-adjusted p<0.01 **(Figure 4 and S4, Table S7.1)**. The top-ranked gene-set was observed in the PTV collapsing analysis model, in all asthma, reflecting genes intolerant to loss-of-function mutations in human populations, according to the gnomAD pLI score[28] (pLI>0.9, n=3230 genes, FDR p=4.1×10^−5^). The second-ranked gene-set is drawn from the same pLI gene-set but excludes pLI genes previously reported in OMIM (pLI>0.9, n=2275 genes, FDR p=0.0005). Only 148 genes from the latter gene-set achieved p<0.1 in the original collapsing analysis **(Table S7.2)**. Both gene-sets were also similarly enriched in the early asthma group, though not significantly.

**Figure 4.**
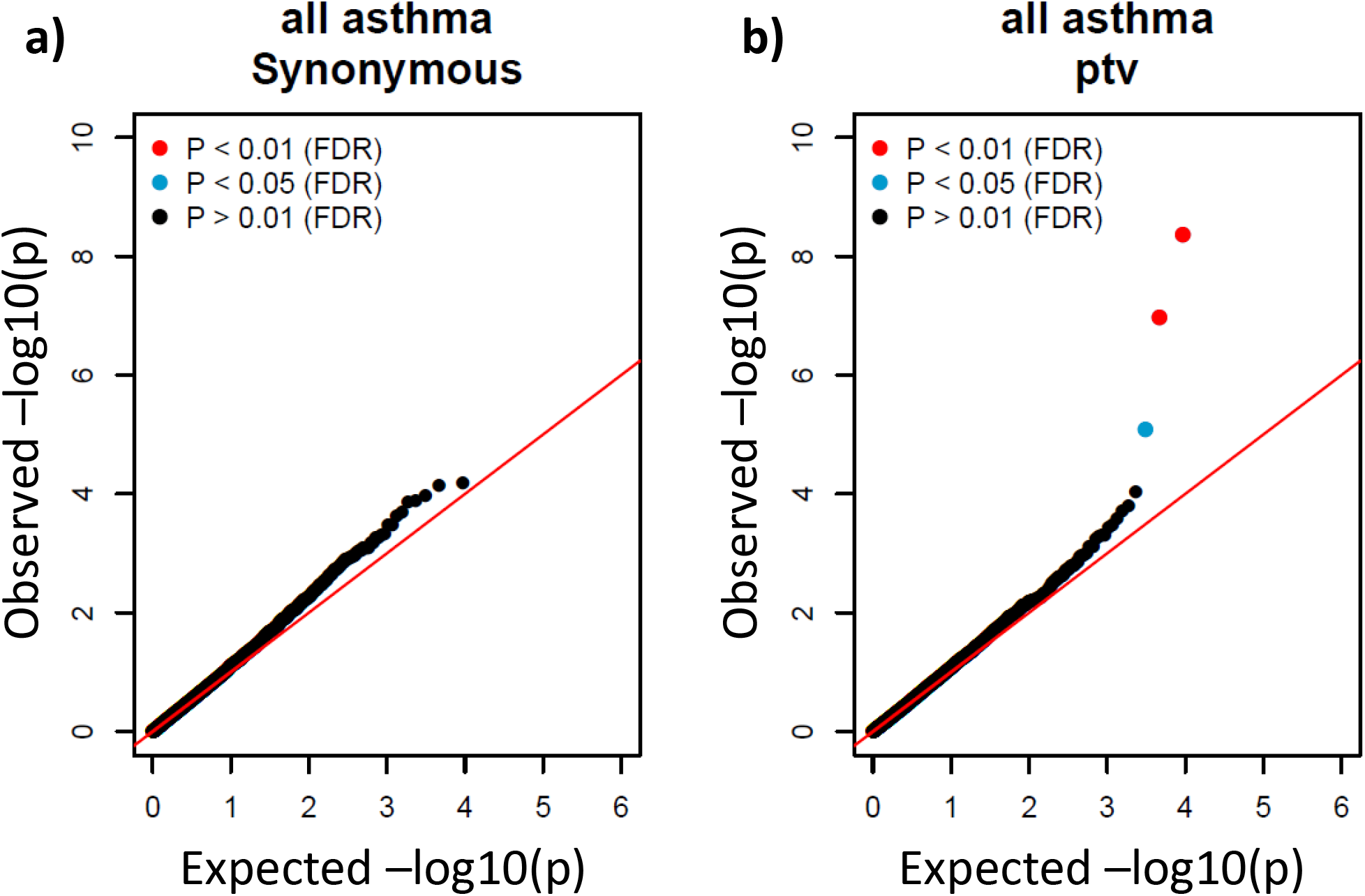
Quantile-quantile plots show the distribution of gene sets in the MegaGene analysis using variants classed as **(A)** rare synonymous variants or **(B)** rare protein-truncating variants and all asthma cases.

## Discussion

This work demonstrates the value of exome sequencing to gain insight into the genetic architecture of asthma, even among known genes. Within *FLG*, two well-described PTVs (rs558269137-CACTG-C NP_002007.1:p.Ser761fs and rs61816761-G-A NP_002007.1:p.Arg501Ter) have each independently risen to relatively common frequencies (MAF ~1%) in global populations[28]. In our data, comprising of sequences from 24,576 subjects with asthma, >12% of West Europeans carry at least one *FLG* PTV, including hundreds of rare-to-private PTVs. Exome sequence data allowed us to further catalogue the full allelic spectrum of genetic variation in *FLG* and establish with high confidence that the effect of rare and ultra-rare *FLG* PTVs is comparable to the two common *FLG* PTVs. Moreover, by comparing *FLG* PTV asthma risk across age groups, we show that *FLG* PTVs are most relevant to childhood-onset asthma, with no evidence of an increase in adult-onset asthma in heterozygotes. Further, we confirmed that this increased risk is significant and substantial in those who otherwise report no allergy phenotype. Finally, we observed that the *FLG* PTV genetic model appears to be additive, whereby recessive *FLG* PTV genotypes carry a significantly greater risk of developing asthma compared to single *FLG* PTV carriers.

Common noncoding (presumed regulatory) variants mapped to *IL33* have been previously linked to an increase in *IL33* expression and asthma risk[15]. In contrast to *FLG*, where we observed a PTV class effect, in *IL33* the protective PTV signal currently appears specific to a single, previously reported, essential splice variant[15] (GRCh38 chr9:g.6255967G>C, rs146597587-C)[24]. We observed a significant decrease in eosinophil count in carriers of this splice variant, but no significant association with eosinophils among the collection of remaining *IL33* PTVs. The absence of a similar protective signal among these rarer *IL33* PTVs is puzzling, since rs146597587 was functionally characterised as a loss-of-function variant[24], and two clinically efficacious IL33-depleting therapies were recently reported[29], [30].

Our exWAS analysis replicated known asthma associations across 44 variants in 33 coding genes **(Figure S2)**. Although our exWAS did not observe significant associations with variants in *ORMDL3, IL1RL1*, or *GSDMB*, three genes commonly linked to asthma in microarray GWAS, the exWAS only examines the contribution of protein-coding variants, suggesting these previously reported signals might be driven by noncoding regulatory aberrations rather than linkage to protein-coding variation.

Interestingly, in the MegaGene analysis, we observed an enrichment of PTVs occurring in genes that are intolerant to truncating mutations in humans, which was significant in the all asthma cohort, and similar but non-significant in the early-asthma group. It is particularly notable that we observed the significant enrichment only in PTVs and not in other categories of variants. This finding remains something to validate in future studies.

Although asthma is recognised as a heterogenous disease, current stratification of relevant subgroups is limited mainly to Type 2 (T2) inflammation driven disease. Our analysis of two previously recognised risk loci indicates possible genetic identification of further patient subgroups. The lack of association of *FLG* PTVs with eosinophil count, often used as a surrogate biomarker for T2 inflammation, suggests they act potentially independently and may overlap both T2 and non-T2 driven patients. Furthermore, we found that *FLG* is a risk factor of only childhood asthma, and the IL-33 protective loss-of-function variant is linked to a larger IL-33 drive in early onset asthma. This indicates potential new biomarkers of etiological subgroups, and a precision medicine approach may enable the selection of patients with an IL-33 drive may identify a IL-33 inhibitor responsive asthma sub-population.

This study is, to our knowledge, the largest to attempt a comprehensive search for rare, high-risk variants or genes in asthma. We demonstrate how access to the full allele frequency spectrum via sequencing provides clarity on the underlying contribution to disease risk for previously implicated genes. For other common complex conditions in which the risk contribution of rare variation is unknown, sequencing can help illuminate the genetic architecture. With the advent of large biobanks with comprehensive phenotyping, such as the UKB, such studies will be increasingly possible for a wider range of conditions.

## Acknowledgements

During this work, SCC was supported by the AstraZeneca postdoc program. We would like to thank David B Goldstein from Columbia University Medical Centre for his valuable feedback during the early stages of this research study. We would like to thank all participants and organisers of the UK Biobank for creating an open scientific resource for the research community. This work was referring UKB application 26041. We would also like to thank the investigators involved in the SAGA study, and all participants of the original SAGA study.

