## Supplementary Material for "A broad exome study of the genetic architecture of asthma reveals novel patient subgroups"

#### Section S1-A: Diagnosis and demographics in SAGA

Exome data was available from 422 SAGA study index cases. All SAGA participants, including parents and siblings with no prior diagnosis of asthma, were evaluated using (i) a clinical history questionnaire, (ii) a BHR assessment using a methacholine challenge, (iii) skin prick test to a range of common aero-allergens, (iv) blood sampling for total IgE measurement, and (v) blood sampling for specific IgE measurement by Phadiatop. SAGA probands were diagnosed at a median age of three years old, and 99% reported an onset of asthma before age 18.

#### Section S1-B: Diagnosis and demographics in UK BioBank

Among 200,603 participants of the UKB with sequencing data available at the time of this study we sought to identify participants with a reported history of asthma using the following codes:

- Field 41270 – ICD 10 asthma codes: J45, J450, J451, J458, J459
- Field 41271 – ICD 9 asthma codes: 4930, 4931, 4932, 4938, 4939
- Field 20002 – Self-reported asthma code: 1111
    - For age at diagnosis in UKB, this was restricted to the self-reported participants, where the asthma diagnosis achieved a median age of 33.5.

We adopted the KING relatedness coefficient ≥ 0.125 to ensure our final test cohort did not include subjects with second-degree or closer relatives. When a pair of subjects were related up to 3$^{rd}$ degree, the subject with higher average exome coverage was retained to ensure only unrelated index subjects were included in the tests.

#### Section S1-C: Codes for allergic phenotypes from the UKBB subjects

For exclusion of allergic subjects in our *FLG* analysis, we focused on UKB dataset as it provided rich information on participants with allergic phenotypes, either self-reported (20002) or recorded in ICD10 (41270/41202/41204) and ICD9 (41271/41203/41205) records. This included subjects with eczema/dermatitis, psoriasis, allergic rhinitis, unspecified allergy or food allergies, a total of 10.9% of all UKB subjects. We used self-reported data (Code 20002, illness codes: 1452, 1453, 1387, 1374, 1668, 1386, 1385, 1670, 1669, 1671) and the primary and secondary ICD10 (41270/41202/41204 codes: L409, L309, 6961, J310, J301, 4779, J450, L500, L239, L400, J304, L208, J303, J300, 6929, L259, L209, 4778, D690, L231, 6928, L235, L308, J302, 4720, L408, L22, L249, L234) and ICD9 (41271/41203/41205 codes: 69180, 7080, 69180, 6918, 4770).

#### Section S2: Post-hoc exWAS filters

We applied high quality variant calling thresholds to mitigate the effects of low-quality variant calls (Table S2). Following these, we performed additional variant-level QC.

1) We excluded indels that are not already represented among a collection of 125,748 independently exome sequenced subjects from the gnomAD reference cohort.
2) We excluded variants that violated Hardy-Weinberg Equilibrium with an exact test p-value < $2x10^{-5}$ among case or control cohorts.
3) We excluded variants with >1% MAF in gnomAD but with a frequency in our control cohort of <0.05%.

#### Section S3: MegaGene gene set selection

We tested a total of 9341 gene sets for each genetic model. These are standardised and designed to be unbiased or relevant to a wide range of diseases. Gene sets containing two different genes with overlapping CCDS regions were excluded. Gene sets comprise the following:

1. 8390 gene sets from Gene Set Enrichment Analysis (GSEA) [1]
2. 912 gene families from HUGO gene nomenclature committee GNC [2]
3. 33 gene sets associated with various diseases: cancer (https://www.cancer.gov/tcga), chronic kidney disease, epilepsy.
4. 6 gene sets based on loss-of-function evidence, using the human population-based pLI score[3] and mouse gene knockouts from the mouse genome informatics databased [4].

#### Section S4: MegaGene pLI enrichment

To further test the robustness of the top MegaGene results we applied the following additional checks:

1) Individuals recruited from the SAGA trial were whole-genome sequenced at a different facility to UKB exomes. We excluded SAGA case participants and re-ran MegaGene analysis. The p-value in the pLI gene sets remained highly significant ($p=1.7\times10^{-8}$), reassuring us this was not driven by a batch effect.
2) In the pLI genes carrying enrichment signal ($p<0.1$, n=148) we observed that the vast majority (86%) also had an FDR LoF < 0.01[3]. This suggests that the genes driving the observed signal are consistently reported as LoF-depleted using an independent metric to pLI.
3) Further restricting to the most enriched pLI genes ($p<0.01$, n = 13) we reviewed the PTV genotypes driving those enrichment signals and found that they were not driven by individual variants or indels that were significantly different between gnomAD non-Finnish Europeans and our internal European control cohort.

**Commented [PS1]:** need to cite the paper for this score

Figure S1: Distribution of age of onset among 18,533 UKB participants with self-reported asthma.

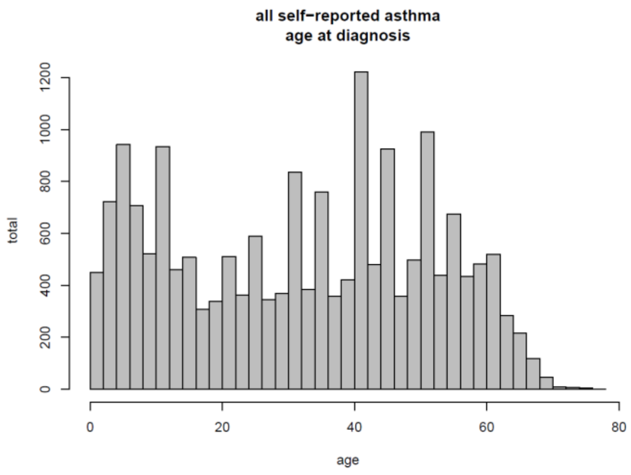

Figure S2: exWAS Manhattan plots

Red line denotes the $p<5x10^{-8}$ threshold. Red gene names indicates the variant was determined to be a likely artefact on manual examination.

#### Figure S2.1 All asthma

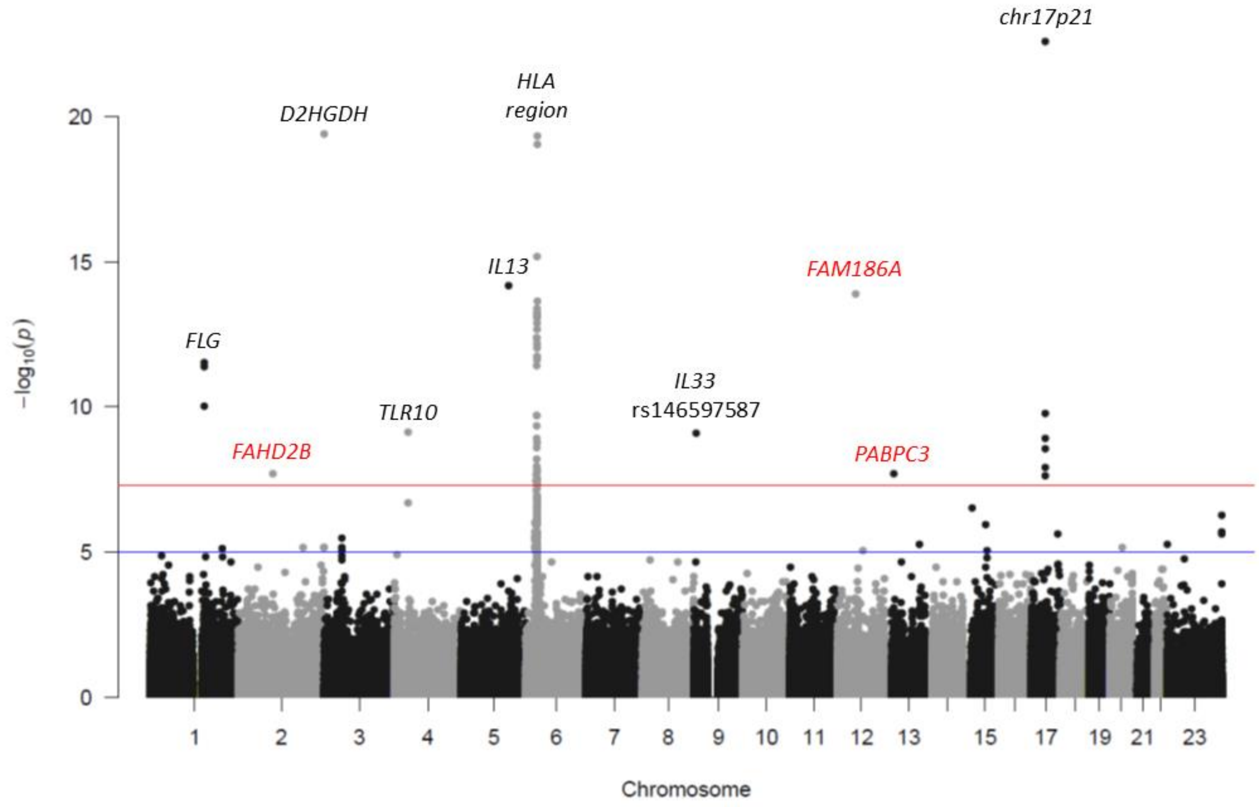

Figure S2.2 Early-onset asthma

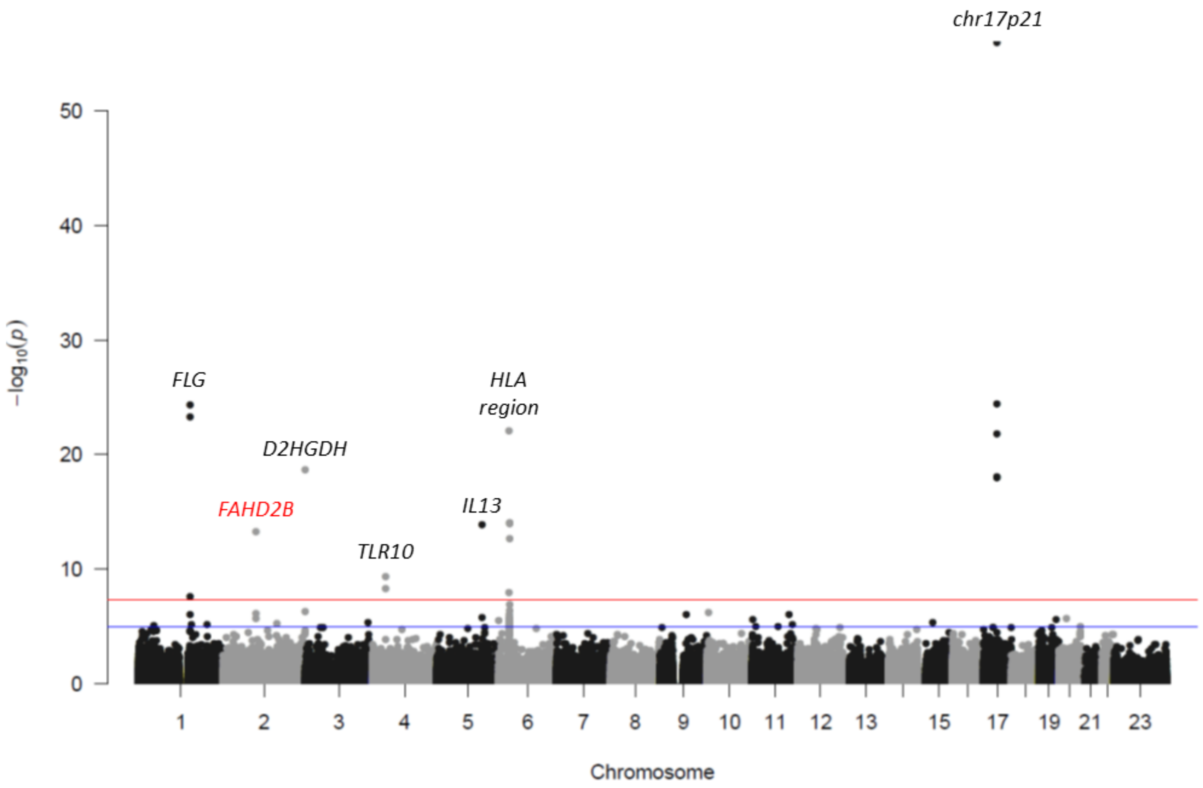

Figure S3: Collapsing analysis QQplots for all asthma

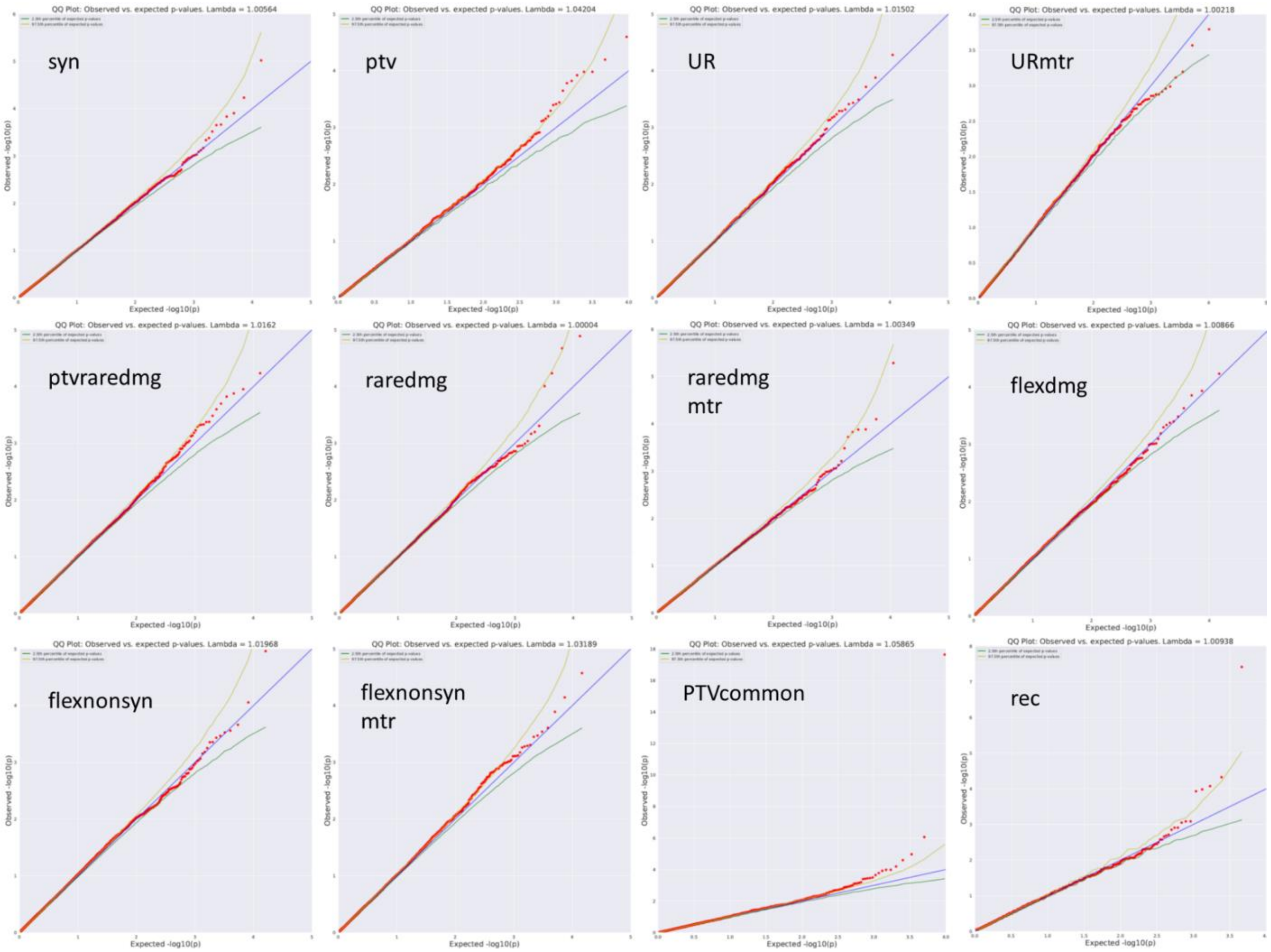

Figure S4: MegaGene QQplots

E.4.1: MegaGene plots from all-asthma group

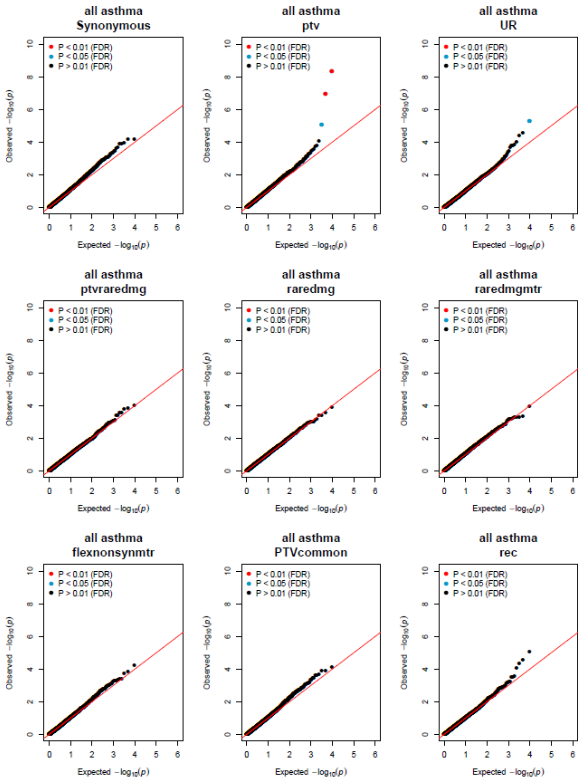

E.4.2: MegaGene plots from early-onset asthma group

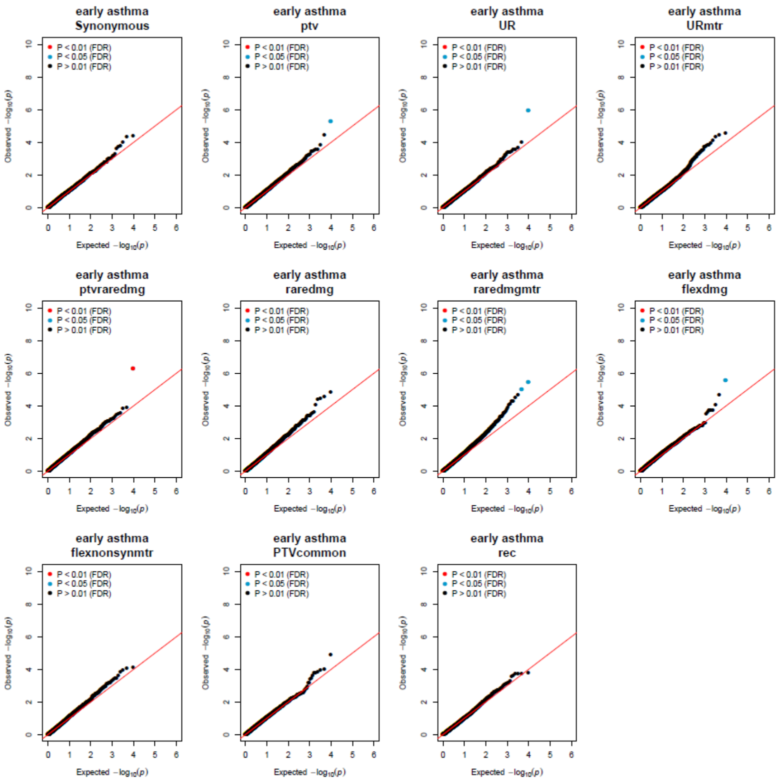

Table S1. Control selection

Case selection used the same QC criteria but did not exclude samples with respiratory conditions or any FEV/FV1 limitations.

| All UKB + SAGA (n) | 201,042 | |
|---|---|---|
| Exclusion criteria | N excluded | % remaining |
| QC | 207 | 99.9 |
| Ancestry non-European | 13234 | 93.3 |
| Asthma diagnosis/reported | 25567 | 80.6 |
| Other Resp conditions | 24622 | 68.4 |
| FEV1/FVC<70% | 16882 | 60.0 |
| Final included controls (n) | 120,530 | |

Table S2. Collapsing analysis variant quality filters

| Argument | Set at | Description of filter |
|---|---|---|
| --min-coverage | 10 | minimum coverage (read depth) to call variants. |
| --ccds-only | NA | Restricts the analysis to variants with annotations in CCDS transcripts only. (currently CCDS r22; November 2018). |
| --het-percent-alt-read | 0.3-0.8 | Excludes heterozygous variants achieving an alternative allele ratio beyond this range. |
| --min-het-binomial-p | 0.000001 | Excludes heterozygous variants that achieve a one-sided binomial exact test p-value below user input; whereby the binom.test(# alt allele reads, # total read depth, p=0.5, alternative="less") < user input p-value. |
| --gq | 30 | minimum Genotype Quality (GQ) score. |
| --snv-fs | 60 | maximum Fisher's Strand Bias (FS) for SNV calls. |
| --indel-fs | 200 | maximum Fisher's Strand Bias (FS) for INDEL calls. |
| --mq | 40 | minimum Mapping Quality (MQ) score. |
| --qual | 30 | minimum QUAL score. |
| --rprs | -2 | minimum read position rank sum (RPRS) score. |
| --mqrs | -8 | minimum mapping quality rank sum (MQRS) score. |
| --var-status | PASS | Classified by Dragen variant caller as a "PASS" variant; requires a quality score of 'DRAGENHardQUAL' < 10.4 |
| --hom-percent-alt-read | 0.8-1.0 | Exclude homozygous genotypes achieving an alternative read frequency outside defined range. |
| --min-covered-sample-binomial-p | 0.000001 | For a given variant site, compare the proportion of cases among the test cohort with user-defined coverage levels to the proportion of cases that make up the test cohort. Exclude variants achieving a binomial exact test p-value below user defined threshold. |
| --max-missing-cohorts-pcnt | 1 | maximum percentage of cases or controls that can have missing genotypes for a variant. Akin to variant missingness of 1%. |
| --gnomad-exome-cov-10x | 0.25 | Proportion of all Gnomad exomes where the read depth at the given variant site exceeds 10x, is greater than or equal to the desired value. |
| --gnomad-exome-z-score | -2.0 | Z-score from Wilcoxon rank sum test of ALT vs. REF read position (RPRS) bias in exomes is greater than or equal to the provided value. |
| --gnomad-exome-mq | 30 | Joint-calling using exome based RMS Mapping Quality is greater than or equal to the provided value. |
| --gnomad-exome-ac-raw-pct | 50 | (Percentage) Only include variants where "gnomAD Exome AC" / "gnomAD Exome AC_raw" is greater than or equal to N (or equals NA) |

Table S3. Collapsing analysis qualifying variant models

**High-impact variants:** exon loss variant, frameshift variant, start lost, stop gained, stop lost, splice acceptor variant, splice donor variant, gene fusion, bidirectional gene fusion, rare amino acid variant, transcript ablation

**Moderate-impact variants:** conservative inframe deletion, conservative inframe insertion, disruptive inframe insertion, disruptive inframe deletion, missense variant splice region variant, missense variant, protein altering variant

**Missense variants:** missense variant splice region variant, missense variant

| Code | Description | Genotype | Predicted Variant Function | Test Cohort MAF | gnomAD maximum global MAF | gnomAD maximum MAF in ancestry group | Maximum sample QC failure (n or %) | Minimum REVEL score (missense) | Minimum MTR score | Minimum MTR intragenic percentile |
|---|---|---|---|---|---|---|---|---|---|---|
| syn | Synonymous | Dominant | Synonymous | 0.0005 | 0.00005 | NA | 2 | 0.25 | NA | NA |
| ptv | Rare, Protein Truncating Variants | Dominant | High impact | 0.001 | 0.001 | 0.001 | 4 | 0.25 | NA | NA |
| UR | Ultra-Rare damaging variants | Dominant | High impact + Moderate impact | 0.00025 | 0 | NA | 1 | 0.25 | NA | NA |
| URmtr | Ultra-Rare damaging variants MTR<50% | Dominant | High impact + Moderate impact | 0.00025 | 0 | NA | 1 | 0.25 | 0.78 | 50% |
| ptvraredmg | Rare, Protein Truncating Variants & Very rare missense | Dominant | High impact + Moderate impact | 0.001 (high) \| 0.0005 (moderate) | 0.001 (high) \| 0.0005 (moderate) | 0.001 (high) \| 0.0005 (moderate) | 4 | 0.25 | NA | NA |
| raredmg | Rare Damaging Missense | Dominant | Missense | 0.0005 | 0.00005 | NA | 2 | 0.25 | NA | NA |
| raredmgmtr | Rare Damaging Missense MTR<50% | Dominant | Missense | 0.0005 | 0.00005 | NA | 2 | NA | 0.78 | 50% |
| flexdmg | Flexible Damaging variants | Dominant | High impact + Moderate impact | 0.001 | 0.0005 | 0.001 | 4 | 0.25 | NA | NA |
| flexnonsyn | Flexible Non-Synonymous variants | Dominant | High impact + Moderate impact | 0.001 | 0.0005 | 0.001 | 4 | NA | NA | NA |
| flexnonsynmtr | Flexible Non-Synonymous variants MTR<50% | Dominant | High impact + Moderate impact | 0.001 | 0.0005 | NA | 4 | NA | 0.78 | 50% |
| rec | Recessive Rare | Recessive | High impact + Moderate impact | 0.005 | 0.0005 | 0.005 | 2 | NA | NA | NA |
| PTVcommon | Protein Truncating Variants, Common | Dominant | High impact | 0.05 | 0.05 | 0.05 | 0.5% | NA | NA | NA |

#### Table S4: Collapsing Analysis results

[This will be a table of the full collapsing analysis results summary – will be submitted as a separate table]

#### Table S5: ExWAS results

##### S5A: All asthma, allelic exWAS results, $p<5.0\text{x}10^{-08}$

| Gene Name | Chr | Base (hg38) | Ref | Alt | MHC region | Function | Case MAF % | Control MAF % | P | Odds Ratio | Previous GWAS asthma phenotype for gene | Citation (PubMed ID) |
|---|---|---|---|---|---|---|---|---|---|---|---|---|
| *ZPBP2* | 17 | 39868373 | C | T | ·· | synonymous | 45·3 | 47·8 | $2{\cdot}7\text{x}10^{-23}$ | 0·91 | Asthma | 2743920, 2086050, 27182965 |
| *D2HGDH* | 2 | 241751260 | G | A | ·· | missense | 23·7 | 25·7 | $4{\cdot}0\text{x}10^{-20}$ | 0·90 | Asthma | 27182965 |
| *TAP2* | 6 | 32828974 | T | C | Yes | missense | 22·9 | 24·9 | $4{\cdot}5\text{x}10^{-20}$ | 0·90 | ·· | ·· |
| *TAP2* | 6 | 32829520 | T | C | Yes | synonymous | 22·9 | 24·8 | $8{\cdot}8\text{x}10^{-20}$ | 0·90 | ·· | ·· |
| *TNXB* | 6 | 32061449 | A | G | Yes | synonymous | 30·4 | 32·3 | $6{\cdot}5\text{x}10^{-16}$ | 1·09 | Eosinophil counts, atopic dermatitis | 27863252, 23886662, 25574825 |
| *IL13* | 5 | 132660272 | A | G | ·· | missense | 19·3 | 17·8 | $6{\cdot}6\text{x}10^{-15}$ | 0·91 | Asthma | 29273806 |
| *TAP2* | 6 | 32830032 | C | T | Yes | missense | 11·0 | 9·8 | $2{\cdot}2\text{x}10^{-14}$ | 1·13 | ·· | ·· |
| *VWA7* | 6 | 31766568 | G | T | Yes | synonymous | 14·3 | 13·0 | $4{\cdot}0\text{x}10^{-14}$ | 1·11 | ·· | ·· |
| *TNXB* | 6 | 32067917 | C | T | Yes | synonymous | 14·1 | 12·9 | $5{\cdot}4\text{x}10^{-14}$ | 1·11 | Eosinophil counts, atopic dermatitis | 27863252, 23886662, 25574825 |
| *TNXB* | 6 | 32061428 | G | A | Yes | synonymous | 14·1 | 12·9 | $6{\cdot}4\text{x}10^{-14}$ | 1·11 | Eosinophil counts, atopic dermatitis | 27863252, 23886662, 25574825 |
| *VWA7* | 6 | 31765689 | T | C | Yes | missense | 14·3 | 13·0 | $7{\cdot}1\text{x}10^{-14}$ | 1·11 | ·· | ·· |
| *MSH5* | 6 | 31760120 | C | T | Yes | synonymous | 14·2 | 13·0 | $8{\cdot}6\text{x}10^{-14}$ | 1·11 | ·· | ·· |
| *TNXB* | 6 | 32096949 | T | C | Yes | missense | 14·1 | 12·9 | $1{\cdot}2\text{x}10^{-13}$ | 1·11 | Eosinophil counts, atopic dermatitis | 27863252, 23886662, 25574825 |
| *NOTCH4* | 6 | 32201368 | G | A | Yes | synonymous | 6·6 | 5·7 | $2{\cdot}0\text{x}10^{-13}$ | 1·16 | Asthma | 21804548 |
| *MICB* | 6 | 31506180 | C | G | Yes | missense | 15·3 | 14·0 | $4{\cdot}4\text{x}10^{-13}$ | 1·11 | Asthma | 29273806 |
| *TNXB* | 6 | 32058086 | C | T | Yes | synonymous | 18·3 | 16·9 | $6{\cdot}6\text{x}10^{-13}$ | 1·10 | Eosinophil counts, atopic dermatitis | 27863252, 23886662, 25574825 |
| *ATF6B* | 6 | 32128224 | C | A | Yes | 5 prime UTR | 13·9 | 12·7 | $9{\cdot}1\text{x}10^{-13}$ | 1·11 | ·· | ·· |

| | | | | | | | | | | | | |
|---|---|---|---|---|---|---|---|---|---|---|---|---|
| *AGER* | 6 | 32183666 | C | T | Yes | missense | 7·2 | 6·3 | $1·7x10^{-12}$ | 1·15 | FEV/FV1 and COPD | 3080456, 28166213 |
| *EGFL8* | 6 | 32167360 | G | A | Yes | synonymous | 6·6 | 5·8 | $2·2x10^{-12}$ | 1·15 | ·· | ·· |
| ***FLG*** | **1** | **152312600** | **CACTG** | **C** | **··** | **frameshift** | **2·9** | **2·3** | **$2·8x10^{-12}$** | **1·24** | **Childhood asthma** | **31036433** |
| ***FLG*** | **1** | **152313385** | **G** | **A** | **··** | **stop gain** | **16·2** | **15·0** | **$3·6x10^{-12}$** | **1·24** | **Childhood asthma** | **31036433** |
| *C6orf15* | 6 | 31111867 | G | A | Yes | synonymous | 2·8 | 2·3 | $4·0x10^{-12}$ | 1·10 | ·· | ·· |
| *RPTN** | 1 | 152155756 | T | C | ·· | missense | 0·027 | 0 | $8·9x10^{-11}$ | Inf | ·· | ·· |
| *ERBB2* | 17 | 39727784 | C | G | ·· | missense | 33·5 | 32·0 | $1·6x10^{-10}$ | 0·94 | Asthma | 2086050, 29273806 |
| *MICB* | 6 | 31505769 | A | G | Yes | missense | 22·1 | 23·4 | $1·8x10^{-10}$ | 1·08 | Asthma | 29273806 |
| *MICA* | 6 | 31411200 | G | A | Yes | missense | 24·5 | 25·9 | $1·9x10^{-10}$ | 0·93 | Asthma | 27182965 |
| *POU5F1* | 6 | 31166166 | C | A | Yes | 5 prime UTR | 30·7 | 32·1 | $4·4x10^{-10}$ | 1·07 | Psoriasis | ·· |
| *TLR10* | 4 | 38773419 | A | G | ·· | synonymous | 0·3 | 0·4 | $7·2x10^{-10}$ | 0·92 | Asthma / Allergy | 29785011 |
| ***IL33*** | **9** | **6255967** | **G** | **C** | **··** | **splice acceptor** | **17·0** | **18·2** | **$8·0x10^{-10}$** | **0·58** | **Asthma** | **29273806** |
| *DPCR1* | 6 | 30952347 | G | A | Yes | missense | 34·5 | 33·0 | $1·1x10^{-09}$ | 1·08 | ·· | ·· |
| *STARD3* | 17 | 39657827 | G | A | ·· | missense | 17·2 | 16·0 | $1·2x10^{-09}$ | 0·94 | Asthma | 30661310 |
| *CFB* | 6 | 31947158 | A | G | Yes | synonymous | 18·5 | 19·6 | $1·5x10^{-09}$ | 1·08 | ·· | ·· |
| *DPCR1* | 6 | 30951614 | G | A | Yes | synonymous | 27·7 | 26·4 | $2·3x10^{-09}$ | 1·08 | ·· | ·· |
| *TCAP* | 17 | 39666058 | A | C | ·· | synonymous | 17·1 | 16·1 | $2·5x10^{-09}$ | 0·94 | Asthma | 30661310 |
| *MICB* | 6 | 31505784 | A | G | Yes | missense | 31·8 | 30·5 | $6·0x10^{-09}$ | 1·06 | Asthma | 29273806 |
| *MICB* | 6 | 31506223 | G | A | Yes | missense | 44·5 | 45·9 | $1·1x10^{-08}$ | 1·06 | Asthma | 29273806 |
| *PSMD3* | 17 | 39981111 | G | A | ·· | synonymous | 31·8 | 30·5 | $1·1x10^{-08}$ | 0·94 | Asthma | 30661310 |
| *SKIV2L* | 6 | 31967790 | G | A | Yes | missense | 2·7 | 3·2 | $1·3x10^{-08}$ | 0·84 | ·· | ·· |
| *AIF1* | 6 | 31616064 | C | G | Yes | 5 prime UTR | 6·6 | 5·9 | $1·7x10^{-08}$ | 1·12 | ·· | ·· |
| *TRIM31* | 6 | 30112719 | G | A | Yes | synonymous | 36·4 | 37·7 | $2·1x10^{-08}$ | 1·08 | Eosinophil percentage | 27863252 |
| *MED24* | 17 | 40023239 | A | G | ·· | synonymous | 18·1 | 17·1 | $2·3x10^{-08}$ | 0·94 | Asthma | 20860503 |
| *HLA-DQB1* | 6 | 32661480 | T | G | Yes | non coding transcript exon | 2·7 | 3·2 | $2·8x10^{-08}$ | 0·85 | Asthma | 20860503, 21804548, 30929738 |
| *UBD* | 6 | 29556180 | A | G | Yes | synonymous | 13·1 | 12·2 | $3·4x10^{-08}$ | 1·09 | ·· | ·· |
| *EGFL8* | 6 | 32166733 | G | A | Yes | missense | 14·1 | 15·0 | $4·2x10^{-08}$ | 1·07 | ·· | ·· |
| *VARS* | 6 | 31781043 | G | A | Yes | synonymous | 24·4 | 23·2 | $4·4x10^{-08}$ | 0·93 | ·· | ·· |

**Protein-truncating variants are in bold.** *Appears to be in linkage with FLG variant

S5B: All asthma, allelic exWAS results, $p<1.0x10^{-05}$

[This will be a table of the ExWAS analysis results – will be submitted as a separate table]

S5C: Early asthma, allelic exWAS results, $p<1.0x10^{-05}$

[This will be a table of the ExWAS analysis results – will be submitted as a separate table]

Table S6: Comparison of odds ratios in all-asthma and early-asthma exwas

| Variant ID | Gene Name | P Value all_asthma | Odds Ratio all_asthma | P Value early_asthma | Odds Ratio early_asthma | OR further from 1.0 |
|---|---|---|---|---|---|---|
| 17-39868373-C-T | *ZPBP2* | 2.72E-23 | 0.9058 | 1.10E-56 | 0.7397 | early_asthma |
| 2-241751260-G-A | *D2HGDH* | 4.07E-20 | 0.8995 | 2.12E-19 | 0.8185 | early_asthma |
| 6-32828974-T-C | *TAP2* | 4.55E-20 | 0.8985 | 7.84E-15 | 0.8401 | early_asthma |
| 6-32829520-T-C | *TAP2* | 8.83E-20 | 0.8991 | 1.08E-14 | 0.8408 | early_asthma |
| 6-32061449-A-G | *TNXB* | 6.59E-16 | 1.0902 | 6.16E-05 | 1.0847 | all_asthma |
| 5-132660272-A-G | *IL13* | 6.64E-15 | 0.9056 | 1.23E-14 | 0.833 | early_asthma |
| 6-32830032-C-T | *TAP2* | 2.28E-14 | 1.1311 | 4.42E-06 | 1.1495 | early_asthma |
| 6-31766568-G-T | *VWA7* | 4.01E-14 | 1.1148 | 3.08E-06 | 1.1345 | early_asthma |
| 6-32067917-C-T | *TNXB* | 5.40E-14 | 1.1145 | 8.08E-07 | 1.1431 | early_asthma |
| 6-32061428-G-A | *TNXB* | 6.43E-14 | 1.1143 | 1.83E-06 | 1.1384 | early_asthma |
| 6-31765689-T-C | *VWA7* | 7.18E-14 | 1.1137 | 3.97E-06 | 1.1332 | early_asthma |
| 6-31760120-C-T | *MSH5* | 8.62E-14 | 1.1133 | 3.20E-06 | 1.1347 | early_asthma |
| 6-32096949-T-C | *TNXB* | 1.23E-13 | 1.113 | 1.48E-06 | 1.1397 | early_asthma |
| 6-32201368-G-A | *NOTCH4* | 2.02E-13 | 1.1621 | 7.98E-06 | 1.1866 | early_asthma |
| 6-31506180-C-G | *MICB* | 4.44E-13 | 1.1066 | 0.00026 | 1.1017 | all_asthma |
| 6-32058086-C-T | *TNXB* | 6.67E-13 | 1.0977 | 2.27E-05 | 1.1091 | early_asthma |
| 6-32183666-C-T | *AGER* | 1.74E-12 | 1.1483 | 0.000141 | 1.1514 | early_asthma |
| 6-32167360-G-A | *EGFL8* | 2.28E-12 | 1.1536 | 3.44E-05 | 1.1718 | early_asthma |
| 1-152312600-CACTG-C | *FLG* | 2.83E-12 | 1.2409 | 4.94E-25 | 1.7234 | early_asthma |
| 6-31111867-G-A | *C6orf15* | 3.65E-12 | 1.0993 | 0.0015 | 1.0852 | all_asthma |
| 1-152313385-G-A | *FLG* | 4.00E-12 | 1.2421 | 4.99E-24 | 1.7147 | early_asthma |
| 6-31505769-A-G | *MICB* | 1.89E-10 | 1.0784 | 0.0251 | 1.0516 | all_asthma |
| 6-31411200-G-A | *MICA* | 1.98E-10 | 0.9297 | 6.10E-06 | 0.9059 | early_asthma |
| 6-31166166-C-A | *POU5F1* | 4.47E-10 | 1.0691 | 0.000311 | 1.0763 | early_asthma |
| 4-38773419-A-G | *TLR10* | 7.24E-10 | 0.9226 | 4.36E-10 | 0.8542 | early_asthma |
| 9-6255967-G-C | *IL33* | 8.01E-10 | 0.5799 | 1.06E-05 | 0.4359 | early_asthma |
| 6-30952347-G-A | *DPCR1* | 1.17E-09 | 1.0841 | 0.0222 | 1.0593 | all_asthma |
| 17-39657827-G-A | *STARD3* | 1.20E-09 | 0.9383 | 3.47E-25 | 0.8163 | early_asthma |
| 6-31947158-A-G | *CFB* | 1.58E-09 | 1.0794 | 0.0029 | 1.0744 | all_asthma |
| 6-30951614-G-A | *DPCR1* | 2.39E-09 | 1.0825 | 0.0246 | 1.0584 | all_asthma |
| 17-39666058-A-C | *TCAP* | 2.59E-09 | 0.9358 | 1.57E-22 | 0.817 | early_asthma |
| 6-31505784-A-G | *MICB* | 6.01E-09 | 1.0641 | 0.0013 | 1.0675 | early_asthma |
| 6-31506223-G-A | *MICB* | 1.13E-08 | 1.0629 | 0.0017 | 1.0658 | early_asthma |
| 17-39981111-G-A | *PSMD3* | 1.15E-08 | 0.9447 | 7.90E-19 | 0.8453 | early_asthma |
| 6-31616064-C-G | *AIF1* | 1.76E-08 | 1.1214 | 0.0083 | 1.1074 | all_asthma |
| 17-40023239-A-G | *MED24* | 2.37E-08 | 0.9442 | 9.81E-19 | 0.8401 | early_asthma |
| 6-32661480-T-G | *HLA-DQB1* | 2.84E-08 | 0.8475 | 1.89E-13 | 0.6338 | early_asthma |
| 6-32166733-G-A | *EGFL8* | 4.22E-08 | 1.0658 | 0.000598 | 1.0782 | early_asthma |
| 6-31781043-G-A | *VARS* | 4.45E-08 | 0.9256 | 0.000677 | 0.9123 | early_asthma |
| 1-152219980-G-T | *HRNR* | 0.0203 | 1.2605 | 2.48E-08 | 2.3452 | early_asthma |

### Table S7: MegaGene results

#### Table S7.1 - Table of MegaGene results a non.syn $p < 1x10^{-4}$

| Group | Model | Gene Set | X Intercept | non syn | syn | sex | Total Sample Variant (p) | Case Median Qual | Ctrl Median Qual | Enrichment | FDR Adjusted P | Gene Set length |
|---|---|---|---|---|---|---|---|---|---|---|---|---|
| all | ptv | pLI | 0 | 4.36E-09 | 0.001882 | 0.000275 | 0.312381 | 0.392537 | 0.367804 | Case | 4.07E-05 | 3230 |
| all | ptv | pLI_less_Mend_Jan2018 | 0 | 1.08E-07 | 0.000335 | 0.000331 | 0.499224 | 0.274306 | 0.255483 | Case | 0.000502 | 2275 |
| early | ptvraredmg | genefam_910_FKBP_prolyl_isomerases | 4.53E-160 | 5.14E-07 | 0.791542 | 8.02E-79 | 0.047944 | 0.036683 | 0.026038 | Case | 0.004806 | 18 |
| early | UR | REACTOME_PD1_SIGNALING | 0 | 1.15E-06 | 0.308256 | 1.95E-78 | 0.009373 | 0.006593 | 0.002913 | Case | 0.010739 | 18 |
| all | URmtr | pLI | 0 | 1.54E-06 | 0.001348 | 0.000128 | 0.548362 | 1.550364 | 1.506214 | Case | 0.014379 | 3230 |
| early | flexdmg | genefam_910_FKBP_prolyl_isomerases | 7.77E-123 | 2.59E-06 | 0.738395 | 9.42E-78 | 0.01673 | 0.04353 | 0.032395 | Case | 0.024172 | 18 |
| early | raredmgmtr | GO_SOLUTE_PROTON_SYMPORTER_ACTIVITY | 1.04E-296 | 3.73E-06 | 0.575773 | 3.46E-79 | 0.348314 | 0.030334 | 0.021359 | Case | 0.034877 | 26 |
| all | flexdmg | GO_POSITIVE_REGULATION_OF_LONG_TERM_ SYNAPTIC_POTENTIATION | 5.21E-256 | 4.63E-06 | 0.763232 | 0.000182 | 0.063272 | 0.042887 | 0.036703 | Case | 0.043205 | 14 |
| all | UR | pLI | 0 | 5.06E-06 | 0.000879 | 0.000101 | 0.5388 | 2.428928 | 2.376101 | Case | 0.047248 | 3230 |
| early | ptv | GO_CARBOHYDRATE_TRANSMEMBRANE_ TRANSPORT | 0 | 5.32E-06 | 0.004168 | 7.51E-79 | 0.908604 | 0.015177 | 0.009293 | Case | 0.049668 | 25 |
| all | ptv | pLI_99 | 0 | 8.19E-06 | 0.002623 | 0.000301 | 0.661667 | 0.230332 | 0.215814 | Case | 0.025509 | 1803 |
| all | rec | GO_NUCLEOBASE_CONTAINING_COMPOUND_ TRANSMEMBRANE_TRANSPORTER_ACTIVITY | 0 | 8.75E-06 | 0.903213 | 0.000178 | 0.852997 | 0.005506 | 0.003559 | Case | 0.08177 | 31 |
| early | raredmgmtr | GO_CATION_SUGAR_SYMPORTER_ACTIVITY | 4.35E-297 | 9.55E-06 | 0.551921 | 4.31E-79 | 0.366947 | 0.021402 | 0.014275 | Case | 0.044601 | 15 |
| early | PTVcommon | GO_SAGA_TYPE_COMPLEX | 3.75E-187 | 1.26E-05 | 0.219245 | 1.91E-56 | 0.889493 | 0.012505 | 0.006863 | Case | 0.117793 | 34 |
| early | raredmg | GO_SOLUTE_PROTON_SYMPORTER_ACTIVITY | 2.40E-195 | 1.40E-05 | 0.575598 | 2.59E-79 | 0.063297 | 0.055988 | 0.043905 | Case | 0.08796 | 26 |
| all | URmtr | pLI_99 | 0 | 1.75E-05 | 0.002645 | 0.000136 | 0.99169 | 1.064672 | 1.031331 | Case | 0.07433 | 1803 |
| early | flexdmg | CTCTGGA_MIR520A_MIR525 | 1.95E-122 | 2.17E-05 | 0.982142 | 1.13E-77 | 0.007745 | 0.388346 | 0.356739 | Case | 0.101373 | 158 |
| early | raredmgmtr | GO_GROWTH | 2.18E-285 | 2.18E-05 | 0.23545 | 7.85E-79 | 0.145421 | 0.407819 | 0.374181 | Case | 0.067834 | 410 |
| all | URmtr | pLI_less_Mend_Jan2018 | 0 | 2.39E-05 | 0.000218 | 0.00014 | 0.878641 | 0.982994 | 0.951549 | Case | 0.07433 | 2275 |
| all | UR | pLI_less_Mend_Jan2018 | 0 | 2.59E-05 | 0.000156 | 0.000105 | 0.935704 | 1.499468 | 1.460932 | Case | 0.120901 | 2275 |
| all | rec | GO_NUCLEOSIDE_TRANSPORT | 0 | 2.71E-05 | 0.446682 | 0.000212 | 0.865567 | 0.003141 | 0.001804 | Case | 0.126681 | 16 |
| early | URmtr | GO_REGULATION_OF_EXTENT_OF_CELL_ GROWTH | 0 | 2.87E-05 | 0.562318 | 6.67E-78 | 0.052628 | 0.06535 | 0.051963 | Case | 0.145225 | 101 |
| early | raredmg | genefam_910_FKBP_prolyl_isomerases | 1.14E-195 | 2.89E-05 | 0.83643 | 1.61E-79 | 0.072518 | 0.026725 | 0.019195 | Case | 0.08796 | 18 |
| early | raredmgmtr | GO_REGULATION_OF_MULTICELLULAR_ ORGANISMAL_METABOLIC_PROCESS | 5.74E-298 | 3.18E-05 | 0.167625 | 3.78E-79 | 0.34264 | 0.030502 | 0.022209 | Case | 0.074182 | 38 |
| early | ptv | genefam_1303_Adenosine_deaminases_ acting_on_RNA | 0 | 3.40E-05 | 0.989938 | 1.01E-78 | 0.986754 | 0.006071 | 0.002956 | Case | 0.158968 | 8 |
| early | URmtr | GO_POSITIVE_REGULATION_OF_AXON_EXTENSION | 0 | 3.57E-05 | 0.978581 | 4.40E-78 | 0.038367 | 0.029889 | 0.021605 | Case | 0.145225 | 36 |
| early | raredmg | BIOCARTA_G1_PATHWAY | 9.23E-196 | 3.57E-05 | 0.498853 | 2.08E-79 | 0.066473 | 0.039242 | 0.029889 | Case | 0.08796 | 28 |
| early | raredmg | GO_CATION_SUGAR_SYMPORTER_ACTIVITY | 1.79E-195 | 3.77E-05 | 0.575548 | 2.68E-79 | 0.069493 | 0.038735 | 0.029328 | Case | 0.08796 | 15 |
| all | UR | GO_HISTONE_MONOUBIQUITINATION | 0 | 3.94E-05 | 0.120792 | 0.000132 | 0.163095 | 0.014247 | 0.011077 | Case | 0.122606 | 23 |
| all | flexdmg | GO_LONG_TERM_MEMORY | 2.33E-255 | 4.10E-05 | 0.858196 | 0.000228 | 0.053876 | 0.099419 | 0.090751 | Case | 0.135686 | 28 |
| all | rec | GO_POSITIVE_REGULATION_OF_INTERLEUKIN_ 17_PRODUCTION | 0 | 4.50E-05 | 0.282604 | 0.000196 | 0.844827 | 0.005833 | 0.003966 | Case | 0.140205 | 13 |
| early | URmtr | GO_REGULATION_OF_CELL_SIZE | 0 | 4.66E-05 | 0.294815 | 7.60E-78 | 0.065113 | 0.103006 | 0.085937 | Case | 0.145225 | 172 |
| all | flexdmg | PID_AP1_PATHWAY | 5.27E-255 | 4.95E-05 | 0.43276 | 0.000187 | 0.04837 | 0.150933 | 0.140533 | Case | 0.135686 | 70 |
| early | raredmgmtr | GO_POSITIVE_REGULATION_OF_ MULTICELLULAR_ORGANISMAL_METABOLIC_PROCESS | 3.12E-298 | 5.38E-05 | 0.048382 | 3.54E-79 | 0.365103 | 0.019717 | 0.01345 | Case | 0.084621 | 23 |
| early | raredmgmtr | GO_MEGAKARYOCYTE_DIFFERENTIATION | 9.16E-298 | 5.44E-05 | 0.486778 | 3.23E-79 | 0.372456 | 0.020897 | 0.014484 | Case | 0.084621 | 20 |
| all | flexdmg | GO_REGULATION_OF_DNA_REPLICATION | 4.42E-255 | 5.81E-05 | 0.493891 | 0.00019 | 0.029471 | 0.385276 | 0.368821 | Case | 0.135686 | 161 |
| early | flexnonsynmtr | REACTOME_GPCR_LIGAND_BINDING | 8.27E-110 | 7.27E-05 | 0.498802 | 2.21E-75 | 0.058258 | 0.730962 | 0.781361 | Ctrl | 0.315471 | 408 |
| early | URmtr | REACTOME_ERK_MAPK_TARGETS | 0 | 7.63E-05 | 0.869692 | 5.52E-78 | 0.029023 | 0.010638 | 0.006277 | Case | 0.178237 | 21 |
| all | PTVcommon | GO_PROTEIN_FOLDING | 0 | 7.77E-05 | 0.039356 | 4.90E-05 | 0.268361 | 0.211554 | 0.198344 | Case | 0.38805 | 224 |
| early | raredmgmtr | GO_MEGAKARYOCYTE_DEVELOPMENT | 6.71E-298 | 8.01E-05 | 0.413106 | 3.81E-79 | 0.375917 | 0.019211 | 0.013159 | Case | 0.103591 | 16 |

| | | | | | | | | | | | | | |
|---|---|---|---|---|---|---|---|---|---|---|---|---|---|
| early | flexdmg | GO_RIBONUCLEOPROTEIN_COMPLEX_ DISASSEMBLY | 1.45E-122 | 8.54E-05 | 0.303251 | 6.97E-78 | 0.018615 | 0.03359 | 0.025628 | Case | | 0.266006 | 11 |
| all | rec | GO_RESPONSE_TO_GROWTH_FACTOR | 0 | 8.86E-05 | 0.19778 | 0.000314 | 0.634524 | 0.075335 | 0.083574 | Ctrl | | 0.206814 | 475 |
| early | raredmgmtr | GO_DEVELOPMENTAL_GROWTH | 7.85E-289 | 8.87E-05 | 0.69231 | 5.82E-79 | 0.169361 | 0.345298 | 0.316939 | Case | | 0.103591 | 333 |
| early | flexnonsynmtr | REACTOME_LATE_PHASE_OF_HIV_LIFE_ CYCLE | 1.35E-110 | 8.92E-05 | 0.924586 | 1.95E-75 | 0.027537 | 0.185978 | 0.210743 | Ctrl | | 0.315471 | 104 |
| early | raredmg | AHRARNT_01 | 7.52E-194 | 8.99E-05 | 0.38337 | 2.14E-79 | 0.043557 | 0.215832 | 0.19427 | Case | | 0.167928 | 140 |
| all | ptv | GO_REGULATED_EXOCYTOSIS | 0 | 9.18E-05 | 0.493738 | 0.000395 | 0.885612 | 0.075892 | 0.068638 | Case | | 0.214335 | 224 |
| early | PTVcommon | genefam_1059_SAGA_complex | 8.77E-187 | 9.20E-05 | 0.911549 | 2.67E-56 | 0.915801 | 0.009873 | 0.005354 | Case | | 0.254835 | 17 |
| early | UR | FREAC2_01 | 0 | 9.24E-05 | 0.34953 | 4.02E-78 | 0.027838 | 0.184954 | 0.161911 | Case | | 0.427161 | 260 |

table S7.2 - Table of pLI_Less_Mend genes at p<0.1

| Rank in original collapsing analysis | Chrom | Gene Name | Qual Cases | Unqual Cases | Qual Case Freq | Qual Ctrls | Unqual Ctrls | Qual Ctrl Freq | Enriched Direction | Fet P | Odds Ratio | Odds Ratio LCI | Odds Ratio UCI |
|---|---|---|---|---|---|---|---|---|---|---|---|---|---|
| 3 | 12 | BAZ2A | 7 | 24569 | 2.85E-04 | 2 | 120528 | 1.66E-05 | case | 1.04E-04 | 17.1699 | 3.5666 | 82.6583 |
| 5 | 18 | ANKRD12 | 31 | 24545 | 0.0013 | 61 | 120469 | 5.06E-04 | case | 1.20E-04 | 2.4943 | 1.6184 | 3.8441 |
| 11 | 2 | SEMA4C | 10 | 24566 | 4.07E-04 | 9 | 120521 | 7.47E-05 | case | 4.02E-04 | 5.4511 | 2.2148 | 13.4165 |
| 14 | 10 | CCDC6 | 7 | 24569 | 2.85E-04 | 4 | 120526 | 3.32E-05 | case | 6.96E-04 | 8.5848 | 2.5129 | 29.3287 |
| 19 | 7 | THSD7A | 1 | 24575 | 4.07E-05 | 51 | 120479 | 4.23E-04 | ctrl | 0.0013 | 0.0961 | 0.0133 | 0.6956 |
| 25 | 13 | FLT1 | 10 | 24566 | 4.07E-04 | 12 | 120518 | 9.96E-05 | case | 0.0017 | 4.0882 | 1.7661 | 9.4636 |
| 30 | 15 | DPP8 | 11 | 24565 | 4.48E-04 | 15 | 120515 | 1.25E-04 | case | 0.0021 | 3.5977 | 1.6522 | 7.8339 |
| 32 | X | RBBP7 | 5 | 24571 | 2.04E-04 | 2 | 120528 | 1.66E-05 | case | 0.0022 | 12.2632 | 2.379 | 63.2134 |
| 51 | 17 | NUP85 | 8 | 24568 | 3.26E-04 | 9 | 120521 | 7.47E-05 | case | 0.0038 | 4.3605 | 1.6822 | 11.3032 |
| 57 | 2 | CTNNA2 | 7 | 24569 | 2.85E-04 | 7 | 120523 | 5.81E-05 | case | 0.0045 | 4.9055 | 1.7205 | 13.9867 |
| 75 | 11 | NUMA1 | 11 | 24565 | 4.48E-04 | 18 | 120512 | 1.49E-04 | case | 0.0057 | 2.998 | 1.4158 | 6.3483 |
| 86 | 1 | PRRC2C | 9 | 24567 | 3.66E-04 | 13 | 120517 | 1.08E-04 | case | 0.0068 | 3.3962 | 1.4516 | 7.9461 |
| 121 | 7 | HERPUD2 | 5 | 24571 | 2.04E-04 | 4 | 120526 | 3.32E-05 | case | 0.0096 | 6.1315 | 1.6464 | 22.8356 |
| 137 | 20 | NCOA3 | 19 | 24557 | 7.73E-04 | 45 | 120485 | 3.73E-04 | case | 0.0113 | 2.0716 | 1.2115 | 3.5421 |
| 167 | 6 | UBR2 | 14 | 24562 | 5.70E-04 | 30 | 120500 | 2.49E-04 | case | 0.0143 | 2.2894 | 1.2138 | 4.3182 |
| 180 | 9 | DENND1A | 20 | 24556 | 8.14E-04 | 50 | 120480 | 4.15E-04 | case | 0.0155 | 1.9625 | 1.1682 | 3.2969 |
| 183 | 16 | TAOK2 | 16 | 24560 | 6.51E-04 | 37 | 120493 | 3.07E-04 | case | 0.0159 | 2.1215 | 1.18 | 3.8143 |
| 197 | 6 | HDAC2 | 5 | 24571 | 2.04E-04 | 5 | 120525 | 4.15E-05 | case | 0.0165 | 4.9052 | 1.4199 | 16.9451 |
| 205 | 1 | ELAVL4 | 3 | 24573 | 1.22E-04 | 1 | 120529 | 8.30E-06 | case | 0.017 | 14.7148 | 1.5305 | 141.473 |
| 215 | 7 | STX1A | 3 | 24573 | 1.22E-04 | 1 | 120529 | 8.30E-06 | case | 0.017 | 14.7148 | 1.5305 | 141.473 |
| 221 | 6 | HSP90AB1 | 8 | 24568 | 3.26E-04 | 13 | 120517 | 1.08E-04 | case | 0.0171 | 3.0187 | 1.2511 | 7.2841 |
| 222 | 11 | ST5 | 8 | 24568 | 3.26E-04 | 13 | 120517 | 1.08E-04 | case | 0.0171 | 3.0187 | 1.2511 | 7.2841 |
| 224 | 22 | ZDHHC8 | 8 | 24568 | 3.26E-04 | 13 | 120517 | 1.08E-04 | case | 0.0171 | 3.0187 | 1.2511 | 7.2841 |
| 233 | X | ATG4A | 4 | 24572 | 1.63E-04 | 3 | 120527 | 2.49E-05 | case | 0.0187 | 6.5401 | 1.4636 | 29.2239 |
| 235 | 14 | PSMC1 | 4 | 24572 | 1.63E-04 | 3 | 120527 | 2.49E-05 | case | 0.0187 | 6.5401 | 1.4636 | 29.2239 |
| 237 | 9 | UHRF2 | 4 | 24572 | 1.63E-04 | 3 | 120527 | 2.49E-05 | case | 0.0187 | 6.5401 | 1.4636 | 29.2239 |
| 238 | 4 | YTHDC1 | 4 | 24572 | 1.63E-04 | 3 | 120527 | 2.49E-05 | case | 0.0187 | 6.5401 | 1.4636 | 29.2239 |
| 261 | 16 | VPS4A | 6 | 24570 | 2.44E-04 | 8 | 120522 | 6.64E-05 | case | 0.0206 | 3.6789 | 1.2764 | 10.604 |
| 265 | 3 | EPHA6 | 19 | 24557 | 7.73E-04 | 48 | 120482 | 3.98E-04 | case | 0.0209 | 1.942 | 1.1415 | 3.3041 |
| 321 | 12 | PRPF40B | 10 | 24566 | 4.07E-04 | 20 | 120510 | 1.66E-04 | case | 0.0257 | 2.4528 | 1.148 | 5.2407 |
| 327 | 3 | ISY1-RAB43 | 5 | 24571 | 2.04E-04 | 6 | 120524 | 4.98E-05 | case | 0.0261 | 4.0876 | 1.2474 | 13.395 |
| 332 | 5 | ADAMTS19 | 2 | 24574 | 8.14E-05 | 43 | 120487 | 3.57E-04 | ctrl | 0.0262 | 0.228 | 0.0552 | 0.9414 |
| 348 | 6 | SLC29A1 | 9 | 24567 | 3.66E-04 | 16 | 120514 | 1.33E-04 | case | 0.027 | 2.7594 | 1.2192 | 6.245 |
| 367 | 16 | AP1G1 | 2 | 24574 | 8.14E-05 | 0 | 120530 | 0 | case | 0.0287 | NA | NA | NA |
| 370 | 19 | FKBP8 | 2 | 24574 | 8.14E-05 | 0 | 120530 | 0 | case | 0.0287 | NA | NA | NA |
| 372 | 7 | ING3 | 2 | 24574 | 8.14E-05 | 0 | 120530 | 0 | case | 0.0287 | NA | NA | NA |
| 378 | 14 | SRP54 | 2 | 24574 | 8.14E-05 | 0 | 120530 | 0 | case | 0.0287 | NA | NA | NA |
| 379 | 1 | SSBP3 | 2 | 24574 | 8.14E-05 | 0 | 120530 | 0 | case | 0.0287 | NA | NA | NA |
| 381 | 6 | TFEB | 2 | 24574 | 8.14E-05 | 0 | 120530 | 0 | case | 0.0287 | NA | NA | NA |
| 383 | 8 | VDAC3 | 2 | 24574 | 8.14E-05 | 0 | 120530 | 0 | case | 0.0287 | NA | NA | NA |
| 385 | 5 | ZNF608 | 2 | 24574 | 8.14E-05 | 0 | 120530 | 0 | case | 0.0287 | NA | NA | NA |

| | | | | | | | | | | | | | |
|---|---|---|---|---|---|---|---|---|---|---|---|---|---|
| 392 | 4 | DNAJB14 | 6 | 24570 | 2.44E-04 | 9 | 120521 | 7.47E-05 | case | 0.0295 | 3.2701 | 1.1638 | 9.1883 |
| 394 | 3 | FSTL1 | 6 | 24570 | 2.44E-04 | 9 | 120521 | 7.47E-05 | case | 0.0295 | 3.2701 | 1.1638 | 9.1883 |
| 396 | 14 | IRF2BPL | 6 | 24570 | 2.44E-04 | 9 | 120521 | 7.47E-05 | case | 0.0295 | 3.2701 | 1.1638 | 9.1883 |
| 409 | 2 | OSBPL6 | 7 | 24569 | 2.85E-04 | 12 | 120518 | 9.96E-05 | case | 0.0304 | 2.8614 | 1.1264 | 7.2687 |
| 424 | 7 | CACNA2D1 | 4 | 24572 | 1.63E-04 | 4 | 120526 | 3.32E-05 | case | 0.0324 | 4.905 | 1.2266 | 19.6142 |
| 427 | 1 | CSF1 | 4 | 24572 | 1.63E-04 | 4 | 120526 | 3.32E-05 | case | 0.0324 | 4.905 | 1.2266 | 19.6142 |
| 429 | 11 | DPF2 | 4 | 24572 | 1.63E-04 | 4 | 120526 | 3.32E-05 | case | 0.0324 | 4.905 | 1.2266 | 19.6142 |
| 432 | 17 | METTL16 | 4 | 24572 | 1.63E-04 | 4 | 120526 | 3.32E-05 | case | 0.0324 | 4.905 | 1.2266 | 19.6142 |
| 471 | 12 | SLC4A8 | 9 | 24567 | 3.66E-04 | 18 | 120512 | 1.49E-04 | case | 0.0357 | 2.4527 | 1.1018 | 5.4602 |
| 478 | 19 | CSNK1G2 | 14 | 24562 | 5.70E-04 | 35 | 120495 | 2.90E-04 | case | 0.0362 | 1.9623 | 1.0557 | 3.6476 |
| 489 | 12 | DDN | 0 | 24576 | 0 | 20 | 120510 | 1.66E-04 | ctrl | 0.0368 | 0 | NA | NA |
| 492 | 9 | PTPRD | 0 | 24576 | 0 | 20 | 120510 | 1.66E-04 | ctrl | 0.0368 | 0 | NA | NA |
| 496 | 12 | FGD6 | 4 | 24572 | 1.63E-04 | 56 | 120474 | 4.65E-04 | ctrl | 0.0369 | 0.3502 | 0.127 | 0.9659 |
| 504 | 8 | HOOK3 | 3 | 24573 | 1.22E-04 | 2 | 120528 | 1.66E-05 | case | 0.0371 | 7.3573 | 1.2293 | 44.0347 |
| 505 | 10 | LARP4B | 3 | 24573 | 1.22E-04 | 2 | 120528 | 1.66E-05 | case | 0.0371 | 7.3573 | 1.2293 | 44.0347 |
| 507 | 3 | RBM5 | 3 | 24573 | 1.22E-04 | 2 | 120528 | 1.66E-05 | case | 0.0371 | 7.3573 | 1.2293 | 44.0347 |
| 508 | 19 | RFX2 | 3 | 24573 | 1.22E-04 | 2 | 120528 | 1.66E-05 | case | 0.0371 | 7.3573 | 1.2293 | 44.0347 |
| 519 | 1 | RAP1GAP | 3 | 24573 | 1.22E-04 | 47 | 120483 | 3.90E-04 | ctrl | 0.0372 | 0.313 | 0.0974 | 1.0056 |
| 520 | 1 | ASTN1 | 0 | 24576 | 0 | 21 | 120509 | 1.74E-04 | ctrl | 0.0374 | 0 | NA | NA |
| 540 | 13 | PDS5B | 5 | 24571 | 2.04E-04 | 7 | 120523 | 5.81E-05 | case | 0.0387 | 3.5036 | 1.1119 | 11.0402 |
| 541 | 1 | RGL1 | 5 | 24571 | 2.04E-04 | 7 | 120523 | 5.81E-05 | case | 0.0387 | 3.5036 | 1.1119 | 11.0402 |
| 556 | 1 | ZC3H12A | 0 | 24576 | 0 | 22 | 120508 | 1.83E-04 | ctrl | 0.0398 | 0 | NA | NA |
| 563 | 11 | ATG13 | 6 | 24570 | 2.44E-04 | 10 | 120520 | 8.30E-05 | case | 0.0405 | 2.9431 | 1.0695 | 8.0986 |
| 564 | 2 | CNOT11 | 6 | 24570 | 2.44E-04 | 10 | 120520 | 8.30E-05 | case | 0.0405 | 2.9431 | 1.0695 | 8.0986 |
| 567 | 4 | LCORL | 6 | 24570 | 2.44E-04 | 10 | 120520 | 8.30E-05 | case | 0.0405 | 2.9431 | 1.0695 | 8.0986 |
| 570 | 12 | PLEKHA5 | 6 | 24570 | 2.44E-04 | 10 | 120520 | 8.30E-05 | case | 0.0405 | 2.9431 | 1.0695 | 8.0986 |
| 605 | 11 | NAV2 | 13 | 24563 | 5.29E-04 | 32 | 120498 | 2.66E-04 | case | 0.0443 | 1.9929 | 1.0458 | 3.7977 |
| 623 | 9 | UBAP2 | 25 | 24551 | 0.001 | 77 | 120453 | 6.39E-04 | case | 0.0473 | 1.5929 | 1.0143 | 2.5017 |
| 658 | 11 | PLCB3 | 8 | 24568 | 3.26E-04 | 16 | 120514 | 1.33E-04 | case | 0.0502 | 2.4527 | 1.0495 | 5.7316 |
| 666 | 11 | CKAP5 | 4 | 24572 | 1.63E-04 | 5 | 120525 | 4.15E-05 | case | 0.0506 | 3.924 | 1.0536 | 14.6141 |
| 671 | 17 | LUC7L3 | 4 | 24572 | 1.63E-04 | 5 | 120525 | 4.15E-05 | case | 0.0506 | 3.924 | 1.0536 | 14.6141 |
| 672 | 12 | METAP2 | 4 | 24572 | 1.63E-04 | 5 | 120525 | 4.15E-05 | case | 0.0506 | 3.924 | 1.0536 | 14.6141 |
| 678 | 11 | DSCAML1 | 1 | 24575 | 4.07E-05 | 30 | 120500 | 2.49E-04 | ctrl | 0.0508 | 0.1634 | 0.0223 | 1.1986 |
| 717 | 12 | SFSWAP | 6 | 24570 | 2.44E-04 | 11 | 120519 | 9.13E-05 | case | 0.0539 | 2.6755 | 0.9894 | 7.2353 |
| 733 | 9 | RABGAP1 | 5 | 24571 | 2.04E-04 | 8 | 120522 | 6.64E-05 | case | 0.0543 | 3.0657 | 1.0028 | 9.3719 |
| 734 | 20 | RALGAPB | 5 | 24571 | 2.04E-04 | 8 | 120522 | 6.64E-05 | case | 0.0543 | 3.0657 | 1.0028 | 9.3719 |
| 770 | 13 | TPT1 | 21 | 24555 | 8.55E-04 | 63 | 120467 | 5.23E-04 | case | 0.0576 | 1.6353 | 0.9978 | 2.6803 |
| 778 | 3 | SENP7 | 0 | 24576 | 0 | 18 | 120512 | 1.49E-04 | ctrl | 0.0578 | 0 | NA | NA |
| 785 | 6 | BRD2 | 8 | 24568 | 3.26E-04 | 17 | 120513 | 1.41E-04 | case | 0.0582 | 2.3084 | 0.9961 | 5.3495 |
| 817 | 3 | THOC7 | 12 | 24564 | 4.88E-04 | 30 | 120500 | 2.49E-04 | case | 0.0606 | 1.9622 | 1.0045 | 3.8331 |
| 843 | 17 | ACLY | 10 | 24566 | 4.07E-04 | 24 | 120506 | 1.99E-04 | case | 0.0649 | 2.0439 | 0.9773 | 4.2747 |
| 846 | 9 | ANP32B | 3 | 24573 | 1.22E-04 | 3 | 120527 | 2.49E-05 | case | 0.0649 | 4.9049 | 0.9899 | 24.3034 |
| 847 | 1 | ARID4B | 3 | 24573 | 1.22E-04 | 3 | 120527 | 2.49E-05 | case | 0.0649 | 4.9049 | 0.9899 | 24.3034 |
| 855 | 1 | PTGS2 | 3 | 24573 | 1.22E-04 | 3 | 120527 | 2.49E-05 | case | 0.0649 | 4.9049 | 0.9899 | 24.3034 |
| 865 | 17 | PELP1 | 12 | 24564 | 4.88E-04 | 31 | 120499 | 2.57E-04 | case | 0.0657 | 1.8989 | 0.9751 | 3.698 |
| 880 | 11 | UBASH3B | 8 | 24568 | 3.26E-04 | 18 | 120512 | 1.49E-04 | case | 0.0681 | 2.1801 | 0.9478 | 5.0145 |

| | | | | | | | | | | | | | |
|---|---|---|---|---|---|---|---|---|---|---|---|---|---|
| 890 | 14 | MAP3K9 | 4 | 24572 | 1.63E-04 | 50 | 120480 | 4.15E-04 | ctrl | 0.0684 | 0.3923 | 0.1417 | 1.0862 |
| 892 | 12 | PPP1R12A | 4 | 24572 | 1.63E-04 | 50 | 120480 | 4.15E-04 | ctrl | 0.0684 | 0.3923 | 0.1417 | 1.0862 |
| 916 | 15 | CCPG1 | 12 | 24564 | 4.88E-04 | 32 | 120498 | 2.66E-04 | case | 0.072 | 1.8396 | 0.9474 | 3.572 |
| 936 | 17 | TBX2 | 10 | 24566 | 4.07E-04 | 25 | 120505 | 2.07E-04 | case | 0.0726 | 1.9621 | 0.9423 | 4.0859 |
| 944 | 12 | FKBP4 | 5 | 24571 | 2.04E-04 | 9 | 120521 | 7.47E-05 | case | 0.0731 | 2.725 | 0.9132 | 8.1319 |
| 948 | 17 | NLK | 5 | 24571 | 2.04E-04 | 9 | 120521 | 7.47E-05 | case | 0.0731 | 2.725 | 0.9132 | 8.1319 |
| 950 | 2 | PTPN4 | 5 | 24571 | 2.04E-04 | 9 | 120521 | 7.47E-05 | case | 0.0731 | 2.725 | 0.9132 | 8.1319 |
| 952 | 12 | RFX4 | 5 | 24571 | 2.04E-04 | 9 | 120521 | 7.47E-05 | case | 0.0731 | 2.725 | 0.9132 | 8.1319 |
| 953 | 2 | RRM2 | 5 | 24571 | 2.04E-04 | 9 | 120521 | 7.47E-05 | case | 0.0731 | 2.725 | 0.9132 | 8.1319 |
| 954 | 19 | WIZ | 5 | 24571 | 2.04E-04 | 9 | 120521 | 7.47E-05 | case | 0.0731 | 2.725 | 0.9132 | 8.1319 |
| 955 | 5 | BRIX1 | 4 | 24572 | 1.63E-04 | 6 | 120524 | 4.98E-05 | case | 0.0733 | 3.27 | 0.9227 | 11.5886 |
| 959 | 8 | EBF2 | 4 | 24572 | 1.63E-04 | 6 | 120524 | 4.98E-05 | case | 0.0733 | 3.27 | 0.9227 | 11.5886 |
| 964 | 3 | ISY1 | 4 | 24572 | 1.63E-04 | 6 | 120524 | 4.98E-05 | case | 0.0733 | 3.27 | 0.9227 | 11.5886 |
| 965 | 2 | IWS1 | 4 | 24572 | 1.63E-04 | 6 | 120524 | 4.98E-05 | case | 0.0733 | 3.27 | 0.9227 | 11.5886 |
| 970 | 14 | SIPA1L1 | 4 | 24572 | 1.63E-04 | 6 | 120524 | 4.98E-05 | case | 0.0733 | 3.27 | 0.9227 | 11.5886 |
| 972 | 1 | SNX27 | 4 | 24572 | 1.63E-04 | 6 | 120524 | 4.98E-05 | case | 0.0733 | 3.27 | 0.9227 | 11.5886 |
| 973 | 1 | ZMYM4 | 4 | 24572 | 1.63E-04 | 6 | 120524 | 4.98E-05 | case | 0.0733 | 3.27 | 0.9227 | 11.5886 |
| 975 | 7 | AGAP3 | 1 | 24575 | 4.07E-05 | 27 | 120503 | 2.24E-04 | ctrl | 0.0736 | 0.1816 | 0.0247 | 1.3366 |
| 987 | 21 | TIAM1 | 3 | 24573 | 1.22E-04 | 43 | 120487 | 3.57E-04 | ctrl | 0.0739 | 0.3421 | 0.1061 | 1.1027 |
| 991 | 7 | USP42 | 15 | 24561 | 6.10E-04 | 42 | 120488 | 3.49E-04 | case | 0.0748 | 1.752 | 0.9715 | 3.1597 |
| 1011 | 8 | CNOT7 | 2 | 24574 | 8.14E-05 | 1 | 120529 | 8.30E-06 | case | 0.0763 | 9.8095 | 0.8894 | 108.1899 |
| 1014 | 8 | DPYSL2 | 2 | 24574 | 8.14E-05 | 1 | 120529 | 8.30E-06 | case | 0.0763 | 9.8095 | 0.8894 | 108.1899 |
| 1015 | 10 | EPC1 | 2 | 24574 | 8.14E-05 | 1 | 120529 | 8.30E-06 | case | 0.0763 | 9.8095 | 0.8894 | 108.1899 |
| 1016 | 6 | FBXO5 | 2 | 24574 | 8.14E-05 | 1 | 120529 | 8.30E-06 | case | 0.0763 | 9.8095 | 0.8894 | 108.1899 |
| 1017 | 17 | HELZ | 2 | 24574 | 8.14E-05 | 1 | 120529 | 8.30E-06 | case | 0.0763 | 9.8095 | 0.8894 | 108.1899 |
| 1018 | 7 | JAZF1 | 2 | 24574 | 8.14E-05 | 1 | 120529 | 8.30E-06 | case | 0.0763 | 9.8095 | 0.8894 | 108.1899 |
| 1021 | X | MAOB | 2 | 24574 | 8.14E-05 | 1 | 120529 | 8.30E-06 | case | 0.0763 | 9.8095 | 0.8894 | 108.1899 |
| 1023 | 14 | MTA1 | 2 | 24574 | 8.14E-05 | 1 | 120529 | 8.30E-06 | case | 0.0763 | 9.8095 | 0.8894 | 108.1899 |
| 1027 | 20 | RBM39 | 2 | 24574 | 8.14E-05 | 1 | 120529 | 8.30E-06 | case | 0.0763 | 9.8095 | 0.8894 | 108.1899 |
| 1032 | 22 | TCF20 | 2 | 24574 | 8.14E-05 | 1 | 120529 | 8.30E-06 | case | 0.0763 | 9.8095 | 0.8894 | 108.1899 |
| 1033 | 10 | UPF2 | 2 | 24574 | 8.14E-05 | 1 | 120529 | 8.30E-06 | case | 0.0763 | 9.8095 | 0.8894 | 108.1899 |
| 1034 | 14 | WDR20 | 2 | 24574 | 8.14E-05 | 1 | 120529 | 8.30E-06 | case | 0.0763 | 9.8095 | 0.8894 | 108.1899 |
| 1041 | 15 | SNX1 | 9 | 24567 | 3.66E-04 | 20 | 120510 | 1.66E-04 | case | 0.077 | 2.2074 | 1.005 | 4.8484 |
| 1053 | 16 | ARHGAP17 | 11 | 24565 | 4.48E-04 | 27 | 120503 | 2.24E-04 | case | 0.0789 | 1.9985 | 0.9912 | 4.0294 |
| 1055 | 7 | HDAC9 | 11 | 24565 | 4.48E-04 | 27 | 120503 | 2.24E-04 | case | 0.0789 | 1.9985 | 0.9912 | 4.0294 |
| 1080 | 2 | STARD7 | 7 | 24569 | 2.85E-04 | 15 | 120515 | 1.25E-04 | case | 0.0819 | 2.2891 | 0.9332 | 5.6149 |
| 1099 | 11 | INCENP | 11 | 24565 | 4.48E-04 | 28 | 120502 | 2.32E-04 | case | 0.0832 | 1.9271 | 0.9593 | 3.8714 |
| 1107 | 2 | RAPH1 | 15 | 24561 | 6.10E-04 | 44 | 120486 | 3.65E-04 | case | 0.0842 | 1.6724 | 0.9306 | 3.0054 |
| 1114 | 7 | GIGYF1 | 13 | 24563 | 5.29E-04 | 36 | 120494 | 2.99E-04 | case | 0.0848 | 1.7714 | 0.9394 | 3.3406 |
| 1115 | 1 | PTPRU | 13 | 24563 | 5.29E-04 | 36 | 120494 | 2.99E-04 | case | 0.0848 | 1.7714 | 0.9394 | 3.3406 |
| 1125 | 9 | NUP214 | 18 | 24558 | 7.32E-04 | 55 | 120475 | 4.56E-04 | case | 0.0856 | 1.6055 | 0.9427 | 2.7342 |
| 1180 | 3 | TEX264 | 0 | 24576 | 0 | 15 | 120515 | 1.25E-04 | ctrl | 0.0913 | 0 | NA | NA |
| 1194 | 15 | MGA | 0 | 24576 | 0 | 16 | 120514 | 1.33E-04 | ctrl | 0.0919 | 0 | NA | NA |
| 1211 | 8 | INTS8 | 7 | 24569 | 2.85E-04 | 16 | 120514 | 1.33E-04 | case | 0.0944 | 2.146 | 0.8828 | 5.2169 |
| 1237 | 22 | NIPSNAP1 | 11 | 24565 | 4.48E-04 | 30 | 120500 | 2.49E-04 | case | 0.0965 | 1.7986 | 0.9012 | 3.5895 |
| 1241 | 17 | CHD3 | 0 | 24576 | 0 | 17 | 120513 | 1.41E-04 | ctrl | 0.0966 | 0 | NA | NA |

| | | | | | | | | | | | | | |
|---|---|---|---|---|---|---|---|---|---|---|---|---|---|
| 1242 | 2 | PAPOLG | 0 | 24576 | 0 | 17 | 120513 | 1.41E-04 | ctrl | 0.0966 | 0 | NA | NA |
| 1243 | 6 | PDE7B | 0 | 24576 | 0 | 17 | 120513 | 1.41E-04 | ctrl | 0.0966 | 0 | NA | NA |
| 1246 | 17 | RABEP1 | 0 | 24576 | 0 | 17 | 120513 | 1.41E-04 | ctrl | 0.0966 | 0 | NA | NA |
| 1266 | 8 | OSGIN2 | 9 | 24567 | 3.66E-04 | 23 | 120507 | 1.91E-04 | case | 0.099 | 1.9194 | 0.8881 | 4.1487 |
| 1271 | 6 | UST | 9 | 24567 | 3.66E-04 | 23 | 120507 | 1.91E-04 | case | 0.099 | 1.9194 | 0.8881 | 4.1487 |
| 1283 | 20 | BMP7 | 3 | 24573 | 1.22E-04 | 4 | 120526 | 3.32E-05 | case | 0.0996 | 3.6786 | 0.8232 | 16.4376 |
| 1285 | 4 | CPE | 3 | 24573 | 1.22E-04 | 4 | 120526 | 3.32E-05 | case | 0.0996 | 3.6786 | 0.8232 | 16.4376 |
| 1286 | 5 | CYFIP2 | 3 | 24573 | 1.22E-04 | 4 | 120526 | 3.32E-05 | case | 0.0996 | 3.6786 | 0.8232 | 16.4376 |
| 1289 | 12 | FBXO21 | 3 | 24573 | 1.22E-04 | 4 | 120526 | 3.32E-05 | case | 0.0996 | 3.6786 | 0.8232 | 16.4376 |
| 1290 | 11 | FBXO3 | 3 | 24573 | 1.22E-04 | 4 | 120526 | 3.32E-05 | case | 0.0996 | 3.6786 | 0.8232 | 16.4376 |
| 1292 | 22 | GRAMD4 | 3 | 24573 | 1.22E-04 | 4 | 120526 | 3.32E-05 | case | 0.0996 | 3.6786 | 0.8232 | 16.4376 |
| 1297 | 12 | NAB2 | 3 | 24573 | 1.22E-04 | 4 | 120526 | 3.32E-05 | case | 0.0996 | 3.6786 | 0.8232 | 16.4376 |
| 1307 | 4 | SMARCA5 | 3 | 24573 | 1.22E-04 | 4 | 120526 | 3.32E-05 | case | 0.0996 | 3.6786 | 0.8232 | 16.4376 |
| 1308 | 14 | TRIM9 | 3 | 24573 | 1.22E-04 | 4 | 120526 | 3.32E-05 | case | 0.0996 | 3.6786 | 0.8232 | 16.4376 |
| 1310 | 5 | YIPF5 | 3 | 24573 | 1.22E-04 | 4 | 120526 | 3.32E-05 | case | 0.0996 | 3.6786 | 0.8232 | 16.4376 |
